# Genomic Loci Associated with Grain Protein and Mineral Nutrients concentrations in *Eragrostis tef* under Contrasting Water Regimes

**DOI:** 10.1101/2024.08.06.606859

**Authors:** Muluken Demelie Alemu, Shiran Ben-Zeev, Vered Barak, Yusuf Tutus, Ismail Cakmak, Yehoshua Saranga

## Abstract

Climate change is becoming a global concern, threating agriculture’s capacity to meet the food and nutritional requirements of the growing population. Underutilized crops present an opportunity to address resilience to climate change and nutritional deficiencies. Tef is a stress-resilient cereal crop, producing gluten-free grain of high nutritional quality. However, knowledge is lacking on tef’s diversity of grain nutritional properties, their interaction with environmental conditions (e.g., water availability) and the underlying genomic loci. We assessed the effect of water availability on tef grain nutrient concentrations and identify the associated genomic loci. A collection of 223 tef genotypes, a subset of tef diversity panel 300, were grown in the field under well-watered and water-limited conditions in 2021, and phenotyped for grain protein and mineral concentrations and seed color. A genome-wide association study was conducted using 28,837 single-nucleotide polymorphisms (SNPs) and phenotypic data to identify marker–trait associations (MTAs). Tef grain nutrient concentrations exhibited wide genetic diversity with a significant influence of environment. Protein and most micronutrients were more concentrated under water-limited conditions, whereas most macronutrients were higher in the well-watered environment. A total of 59 SNPs were associated with one or more of the studied traits, resulting in 65 MTAs detected under both water treatments, and providing insights into the genetic basis of grain nutrients. Five SNPs reflected multiple associations, with four detecting the same trait under both treatments (multiple-environment effect), and one associated with both Zn and K (pleiotropic effect). In addition, two pairs of closely linked SNPs reflected multiple-environment effects. While multiple-environment associations provide greater support for the integrity of these MTAs, the pleiotropic locus hints at a common mechanism controlling two mineral ions. The identified MTAs shed new light on the genomic architecture of tef’s nutritional properties and provide the basis to enhance tef grain nutritional quality alongside drought resilience.

## 1 Introduction

With millions of people suffering from malnutrition due to protein and micronutrient deficiencies (FAO, 2020; Stevens et al., 2022), the capacity of agriculture to meet food and nutritional requirements of a growing population in the face of climate change is becoming a global concern (FAO, 2023). Underutilized (orphan) crop species, which are not widely cultivated, present outstanding nutritional value and a high tolerance to abiotic and biotic stresses (Bekkering & Tian, 2019; Cheng, 2018; Hunter et al., 2019). Although they are considered underutilized crops, some are staple crops in their origin and center of diversity. These crops offer the potential to improve food and nutrition security for millions of people (Mabhaudhi et al., 2019; Siddique et al., 2021), but they have not been sufficiently studied or improved (Alemu et al., 2024; Allaby, 2021; Milla & Osborne, 2021; Tadele, 2019).

Tef (*Eragrostis tef* (Zucc.) Trotter) is a C4 cereal crop with a genome size of 622 Mb (VanBuren et al., 2020). Ethiopia is tef’s origin and center of diversity (Vavilov, 1951), where it is the most important food and feed crop (Assefa et al., 2017; Chanyalew et al., 2019; D’Andrea, 2008; Tadele, 2019), and the primary ingredient of injera, the traditional Ethiopian bread (Abewa et al., 2019; Ligaba-Osena et al., 2021). Tef is a prestigious cereal crop with outstanding nutritional value, producing gluten-free grains, rich in minerals (Fe, Cu, Mn, Zn, B, Mo, Na, Ca, Mg, K, P, S), carbohydrates, fat, protein, essential amino acids, vitamins and dietary fiber (Abewa et al., 2019; Ligaba-Osena et al., 2021; Shumoy et al., 2018; Tietel et al., 2020; Villanueva et al., 2022; Zhu, 2018). The concentrations of minerals, carbohydrates, fat, proteins and fiber in tef are comparable to or higher than those in other cereal crops (Hager et al., 2012; Ligaba-Osena et al., 2021; Saturni et al., 2010). Thus, tef is recognized as a most important food crop to combat malnutrition, including mitigation of iron-deficiency anemia, particularly in developing nations (Abewa et al., 2019; Daba, 2017). Its favorable nutritional composition and health benefits have contributed to a global interest in tef as a healthy superfood, especially for gluten-intolerant and sensitive individuals (Assefa et al., 2011; Ligaba-Osena et al., 2021; Tietel et al., 2020; Zhu, 2018).

Tef seed is broadly grouped into white and brown colors (Jifar et al., 2015), varying from ivory white to pale white and light brown to dark brown, respectively (Asefa et al., 2023). Seed color is associated with nutritional properties, where brown tef grain is superior to white grain with respect to most minerals and antioxidants (Abebe et al., 2007; Dame, 2020; Tietel et al., 2020), whereas white grain is superior with respect to essential amino acids and protein (Gebru et al., 2019).

Drought is a primary abiotic stress, significantly limiting crop development and productivity (El Sabagh et al., 2020; Vadez et al., 2024). Drought also influences grain quality, either positively or negatively (Stagnari et al., 2016), by modifying morphological, physiological and biochemical characteristics of crop species (Dietz et al., 2021; El Sabagh et al., 2020). Assessing crop grain quality under different environments and understanding the effects of gene-by-environment interactions on quality traits is crucial for crop improvement. Despite extensive documentation on the effects of drought stress on growth, development and production of various crops, its effects on grain quality have not been sufficiently investigated (Chadalavada et al., 2022; Stagnari et al., 2016). Recently, attention has been drawn to various effects of drought on grain quality of cereal crops using high-throughput phenotyping (Chadalavada et al., 2022; El Sabagh et al., 2020; Kamal et al., 2023). Tef grain nutrient concentrations are influenced by genotype and environmental factors (climatic and edaphic variations) (Abewa et al., 2019), but we are not aware of any prior study on the effect of contrasting water regimes on tef grain quality.

Genome-wide association studies (GWASs) have been used for a range of plant species to dissect complex traits in diversity panels (Atwell et al., 2010; X. Huang & Han, 2014). GWASs have been extensively applied for staple cereal crops, but they are only beginning to be used for underutilized crops (Chapman et al., 2022). Recently, GWASs have been successfully applied to several orphan crops, including tef (Alemu et al., 2024; Woldeyohannes et al., 2022). High-throughput genotyping and phenotyping techniques provide opportunities to examine the genomic loci underlying grain quality (Chadalavada et al., 2022; El Sabagh et al., 2020; Kamal et al., 2023). Although the genomic dissection of grain nutrients in crops remains limited, especially in underutilized crops, grain nutritional quality has been targeted in numerous cereal-breeding programs, including foxtail millet (Jaiswal et al., 2019), finger millet (Puranik et al., 2020), sorghum (Kamal et al., 2023), barley (Nyiraguhirwa et al., 2022), wheat (Liu et al., 2021) and rice (Islam et al., 2022).

Tef genotypes hold a rich gene pool and substantial genetic variation with respect to nutrition, yet this crop remains untapped due to a lack of understanding of its genotypic diversity in nutrient concentrations and bioavailability (Ligaba-Osena et al., 2021), and of the genomic loci underlying grain nutritional properties (Ereful et al., 2022). A better understanding of the diversity in tef grain nutrient concentrations, their interaction with environmental conditions and genomic architecture is crucial for the development of nutritious tef varieties.

We previously reported on the assembly of a tef diversity panel 300 (TDP-300) and its utilization for a GWAS of productivity, phenology, lodging and morphophysiological traits under contrasting water regimes (Alemu et al., 2024). The current follow-up study was targeted to tef grain nutritional properties, including protein, micronutrient and macronutrient concentrations, their responses to water availability and their underlying genomic loci. The combination of our previous (Alemu et al., 2024) and current GWASs shed new light on tef’s responses to water stress and provide solid grounds for further studies and the development of novel tef varieties combining drought resilience and high grain yield and nutritional quality.

## 2 Materials and Methods

### 2.1 Plant materials and genotyping

A wide collection of tef genotypes, collected by the USDA in Ethiopia and maintained in the Israel Plant Gene Bank, was used for the current study. Genotype by sequencing and subsequent data processing and filtering resulted in 28,837 single-nucleotide polymorphism (SNP) markers called across 297 accessions, hereafter termed TDP-300. Further details on genotyping, SNP quality and population structure are presented in Alemu et al. (2024). A subset of 223 genotypes from the TDP-300 grown in the field under well-watered (WW) and water-limited (WL) treatments were phenotyped for grain protein, micronutrients and macronutrients.

### 2.2 Growth conditions and phenotyping

Tef genotypes were grown in 2021, in the field at the Kvutzat Shiller farm in Israel (31.879° N, 34.777° E), under WW (total water applied: 438.3 mm) and WL (total water applied: 211.7 mm) conditions, with three replicates. Soil type at the experimental location was clay, with the upper layer (0–30 cm depth) at sowing time containing 8.1, 18.7 and 60.9 mg/kg soil N, P and K, respectively. Liquid fertilizer was applied via the irrigation system at rates of 52, 30 and 85 kg/ha N, P_2_O_5_ and K_2_O, respectively. No rainfall occurred during the experimental seasons, and average temperatures (min/max) were 16.8/31.3°C. Complete details of the experimental layout, environmental conditions and management are described in Alemu et al. (2024).

#### 2.2.1 Grain protein

Grain protein concentrations were estimated with near-infrared spectrometer (NIRS) DS2500 (FOSS, Denmark). Whole-grain samples were dried at 40°C for 24 h, stored in a sealed plastic bag and cooled to room temperature. Approximately 10 g of each tef grain sample was transferred into a small sample cup (6 cm diameter), scanned, and its NIR spectral reflectance across 1100–2498 nm was recorded at 2-nm intervals, using Mosaic Solo software (FOSS Analytical, Hillerød, Denmark).

A preliminary calibration for the NIRS, based on 51 grain samples of various genotypes and treatments from two experiments conducted in two locations in 2019, was already available. In addition, 74 grain samples were prepared by bulking the three replicates of randomly selected genotypes from both treatments of the current experiment; 33 of these were added to the calibration set (for a total of 84 samples) and the remaining 41 samples were used for validation. N concentrations of the calibration and validation samples were determined at Bactochem’s laboratories (Ness Ziona, Israel) by the Kjeldahl method, using block digestion and steam distillation. Total protein content was then calculated by multiplying the N content by a conversion factor of 6.25. The laboratory protein concentrations of the 84 calibration samples were associated with the respective spectral data using the WinISI 4 software (FOSS Analytical, Hillerød, Denmark). Calibration curve coefficient (R^2^) and cross-validation (1-VR) were 0.99 and 0.98, respectively, whereas the R^2^ of the validation set was 0.90. The calibration models were then applied to the entire set of tef grain samples (3 replicates per genotype under each treatment) to predict their protein concentration.

#### 2.2.2 Grain minerals

Samples for determining tef grain mineral concentrations were prepared by combining 0.5 g of grain from each of the three replicates of a genotype under a specific treatment (WW or WL). The grain samples were first thoroughly washed with tap water and then with deionized water to remove the adhering soil dusts, chaff and other possible contaminants. The washed samples were dried at about 40℃ in a forced-draft oven to a constant weight and then subjected to analyses of the targeted mineral nutrients (Fe, Zn, Cu, Mn, Ca, Mg, K, P, and S). The samples were digested in a closed-vessel microwave system (MarsExpress; CEM Corp; Matthews, NC, USA) in 2 ml of 30% (v/v) premium-grade H_2_O_2_ (Merck, EMSURE®, Darmstadt, Germany) and 5 ml of 65% (v/v) premium-grade HNO_3_ (Merck, EMSURE®, Darmstadt, Germany) as described by Yazici et al., (2021). All dilutions required were performed using ultra-pure water (18.2 MΩ). The measurement of the mineral nutrients in the acid digests was conducted by inductively coupled plasma optical emission spectroscopy (ICP-OES; Agilent 5110 Vertical Dual View). The measurements were verified by using certified standard reference material (SRM) obtained from the National Institute of Standards and Technology (NIST, Gaithersburg, MD, USA).

#### 2.2.3 Grain color

Grain colors were scored visually into two broad categories: white and brown (Figure S1) and used both as an independent variable in the analysis of variance (ANOVA) and a dependent variable in the GWAS.

### 2.3 Statistical analyses

JMP Pro 16.0.0 (SAS Institute Inc., 1989–2021) was used for ANOVA. A full factorial nested ANOVA, including environment (E), seed color (SC), genotypes (G, nested in SC) and interactions, was used to analyze the replicated protein results and calculate the least squares mean values. For the non-replicated (bulk) grain minerals, two-way ANOVA was used to analyze the effects of G and E, whereas G * E interactions were visualized as raincloud plots, using “ggrain” package in R (Allen et al., 2021). The distribution of phenotypic traits was presented by density plot using the “ggplot2” package in R (Wickham et al., 2016). The associations among the tested grain nutrients, as well as grain yield (GY) and thousand seed weight (TSW), in WW and WL environments were assessed using Pearson’s correlation coefficient (Peterson et al., 2014) and principal component analysis (PCA) in R (R Core Team, 2020). All traits were tested for normality using the Shapiro–Wilk test and in the few cases of traits that were not normally distributed, Box–Cox transformation procedure was used (JMP Pro 16.0), resulting in normal distribution of all traits.

### 2.4 Genome-wide associations

Phenotypic and genotypic data (28,837 SNPs) were subjected to GWAS using the most advanced models of the Genome Association and Prediction Integrated Tool (GAPIT3) genetics statistical package (Wang & Zhang, 2021) in R (R Core Team, 2020). A multi-locus GWAS model, Bayesian-Information and Linkage-Disequilibrium Iteratively Nested Keyway (BLINK) (M. Huang et al., 2019), was selected due to the lowest rate of false positives as observed in the QQ plots. The effect of population structure was corrected for by including the kinship matrix (K-model) which was calculated using the VanRaden method (VanRaden, 2008). Principal components (PCs) were not included in the association model due to the low clustering tendency of the population (Alemu et al., 2024). To minimize the rate of false positives, we used the stringent Bonferroni-corrected threshold (0.05/number of SNPs = 1.7E^-06^) (Bonferroni, 1936) to determine significant marker–trait associations (MTAs). Once the association between a certain SNP and a trait met this stringent threshold, a putative MTA was considered if the association of the same SNP with other traits or the same trait under a different environment met a less stringent threshold (*p* < 1E-04).

Neighboring MTAs were considered within the same linkage block (i.e., locus) when the linkage between them (r²) was greater than 0.5 and their physical distance smaller than 97,242 bp, the distance at which linkage disequilibrium (LD) decay decreases to below 0.2 (Remington et al., 2001).

## 3 Results

### 3.1 Tef grain nutrient concentrations

The tested tef genotypes (a subset of TDP-300) exhibited wide variation in grain concentrations of protein, micronutrient (Fe, Zn, Cu and Mn) and macronutrient (Ca, Mg, K, P and S) under both WW and WL treatments (Figure 1), as well as across grain colors (brown and white) (Figure 2).

**Figure 1.**
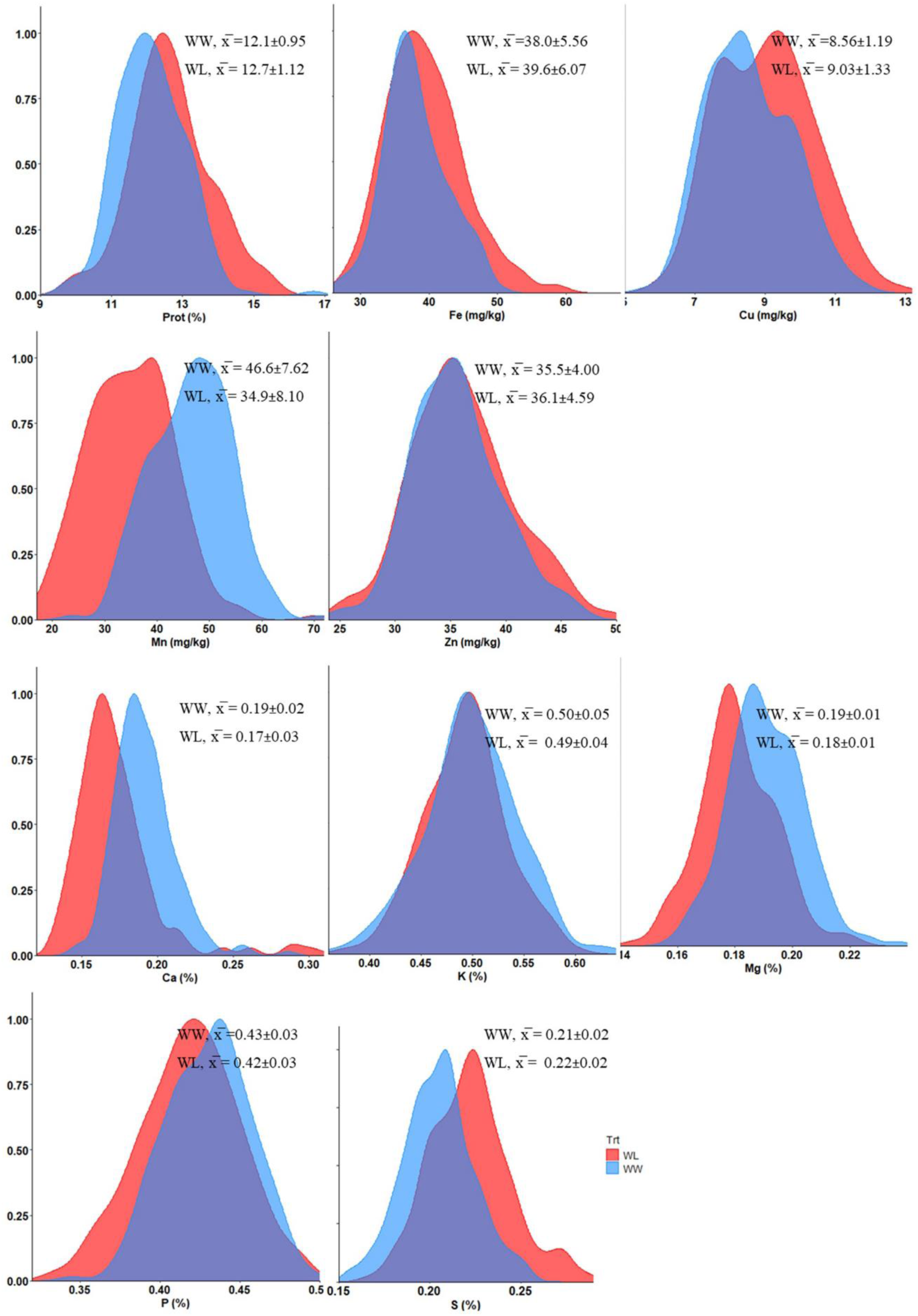
Density plots presenting the distribution of tef grain nutrient concentrations under well-watered (WW, blue) and water-limited (WL, red) treatments in 2021. Prot, protein. ^x̅^ indicates mean values ± SD.

**Figure 2.**
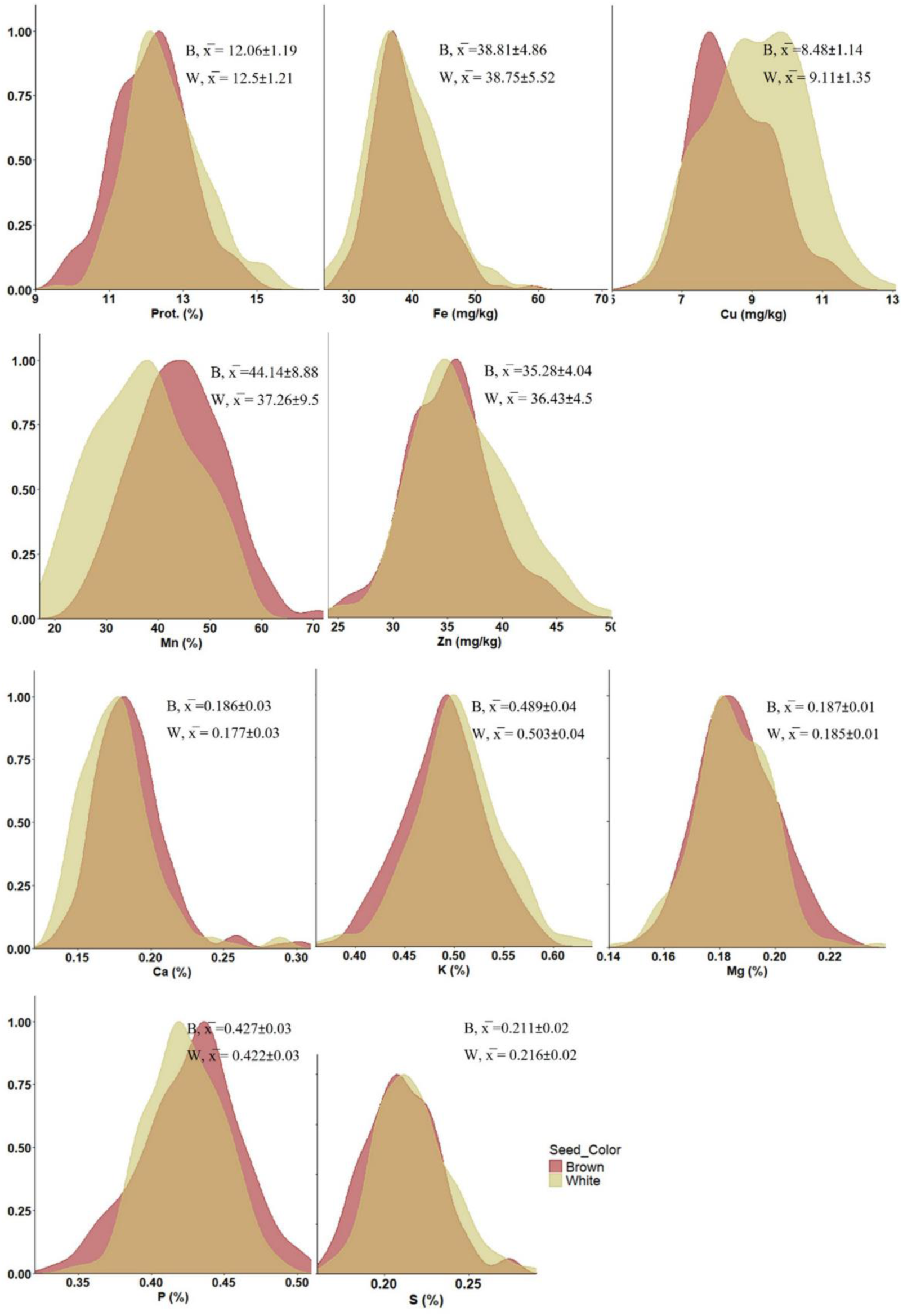
Density plots presenting the distribution of tef grain nutrient concentrations (averaged across treatments) between seed colors: brown (B) (n = 114) and white (W) (n = 109) in 2021. Prot, protein. ^x̅^ indicates mean values ± SD.

Water availability had a highly significant effect (*p* < 0.001) on most mineral nutrients, except Zn which exhibited a lower effect (*p* < 0.05) and K, which did not show any significant effect (Table S1). Grain nutrient concentrations were either positively or negatively affected by water availability (Table S1, Figure 1). Protein and most micronutrients (Fe, Zn and Cu) exhibited higher concentrations under WL vs. WW conditions, whereas Mn exhibited the opposite trend. On the other hand, most macronutrients (Ca, Mg and P) had higher concentrations under the WW vs. WL treatment, except for S which exhibited the opposite trend and K which, as already noted, was not affected. The treatment effects on most nutrient concentrations were in the range of 1–10%, except for Mn which exhibited a 25% lower concentration under the WL vs. WW treatment.

ANOVA revealed significant differences for all grain nutrient concentrations between genotypes (*p* < 0.001), as well as between grain colors (*p* < 0.01), except Fe which did not exhibit significant variation between seed colors (Figure 2, Table S1). Protein, Cu, Zn, K and S exhibited higher concentrations in white seeds. Conversely, Mn, Ca, Mg and P were at higher concentrations in the brown seeds. While most differences in nutrient concentrations between seed colors were in the range of 0–7%, the most pronounced difference was noted for Mn, with an about 18% higher concentration in the brown-seeded vs. white-seeded genotypes.

Treatment-by-genotype interaction, assessed for the replicated protein data by ANOVA, revealed a significant effect (*p* < 0.01), whereas the treatment-by-seed color interaction was not significant (Table S1). For the non-replicated micronutrients and macronutrients, which could not be subjected full factorial ANOVA, raincloud plots (Figure S2) suggested considerable treatment-by-genotype and treatment-by-seed color interactions for most minerals.

### 3.2 Association between phenotypic traits

Correlations among grain nutrients were positive and significant in most cases (Table S2), non-significant in a few cases, and negative and significant in a single case (Mn vs. Cu under WL). In contrast, the correlations between grain yield and grain nutrients were negative in most cases, non-significant in several other cases and positive in three cases (most pronounced for K vs. GY under WW conditions). Finally, the correlations between TSW and grain nutrients were usually non-significant, except three cases of negative correlations (S, Mg and P under WW conditions) and two cases of positive correlation (Fe under both treatments).

PCA, conducted separately for each environment, indicated variations in the components between treatments (Fig. 3). The two main PCs (eigenvalues >1) were extracted, explaining a total of 47.7% and 45.4% of the variation under WW and WL treatments, respectively. PC1 (X-axis) explained 32.5% and 29.8% of the variation and PC2 (Y-axis) explained 15.2% and 15.6% of the variation under WW and WL treatments, respectively. Most of the traits showed positive loadings for PC1 under both the WW and WL treatments, whereas grain yield was negatively loaded in PC1 under both treatments, thus supporting the correlation analyses. Fe and P exhibited the highest contribution to the variation under both treatments. High contributions were also noted for K under WL conditions and for protein under the WW treatment.

**Figure 3.**
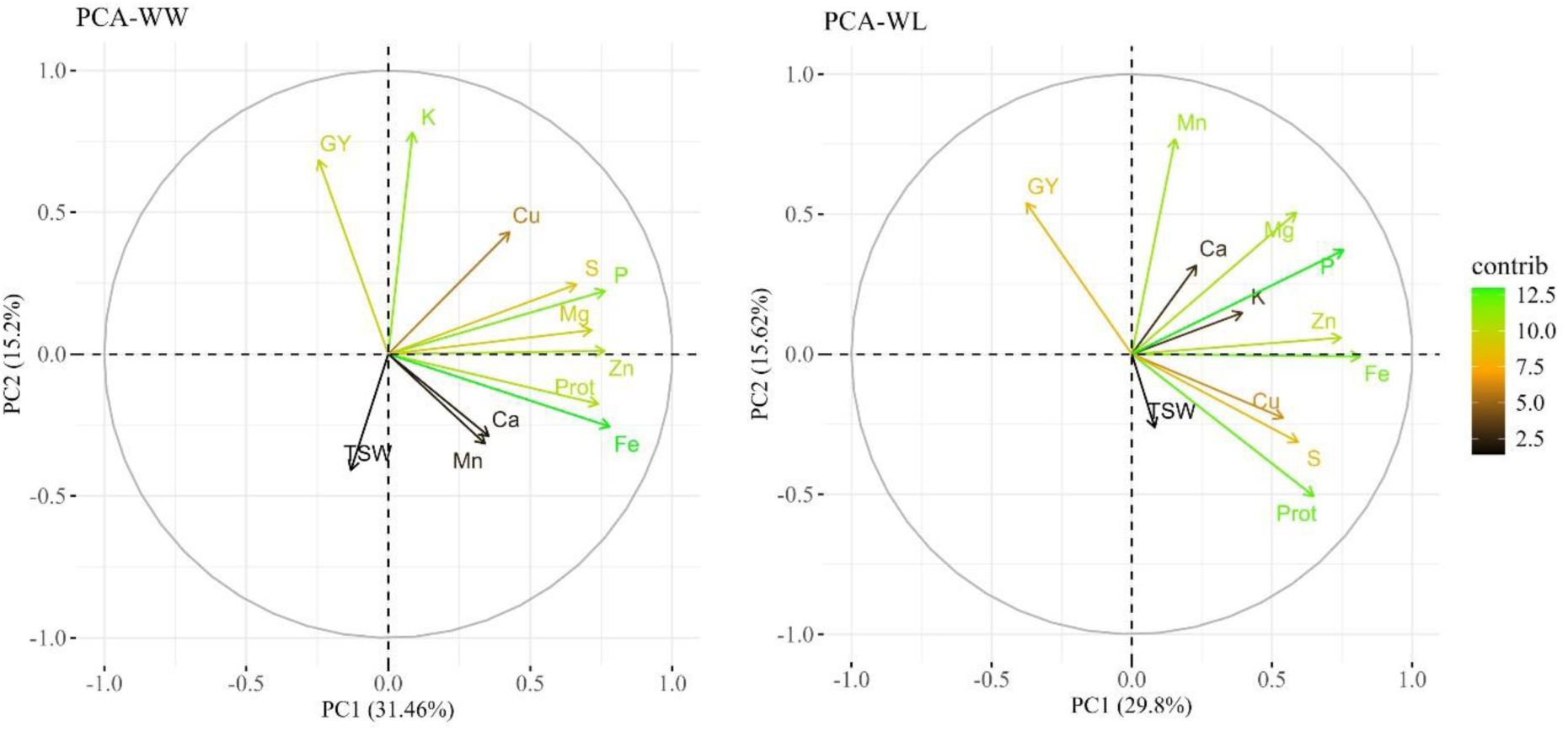
Principal component analysis (PCA) of grain nutrients and yield-related traits of tef diversity panel 300 (TDP-300) under well-watered (WW, left) and water-limited (WL, right) conditions. PCA included grain nutrient concentrations. Prot, protein; GY, grain yield; TSW, thousand seed weight. Biplot vectors showing trait loadings for PC1 and PC2, with color indicating trait contribution to variation.

### 3.3 Genome-wide association

A total of 59 significant MTAs (meeting the Bonferroni-corrected threshold, *p* < 1.7E-06) were identified for seed color and nutrient concentrations in the tef diversity panel (TDP-300) under WW and WL treatments (Tables 1 and 2). An additional four MTAs that met a less stringent threshold (*p* < 1E-04) but overlapped with significant MTAs were also considered and defined as putative MTAs. A greater number of significant MTAs was detected under the WL vs. WW treatment (Table 1). MTAs were unevenly distributed across 19 of the 20 tef chromosomes, with Chr 9B showing no significant association (Figure 4), and their abundance was rather similar on subgenome A (28 SNPs) and subgenome B (31 SNPs). Most tef grain nutritional traits showed significant MTAs under both WW and WL conditions, except for Ca and Zn, which exhibited MTAs only for the WL treatment (Table 1).

**Figure 4.**
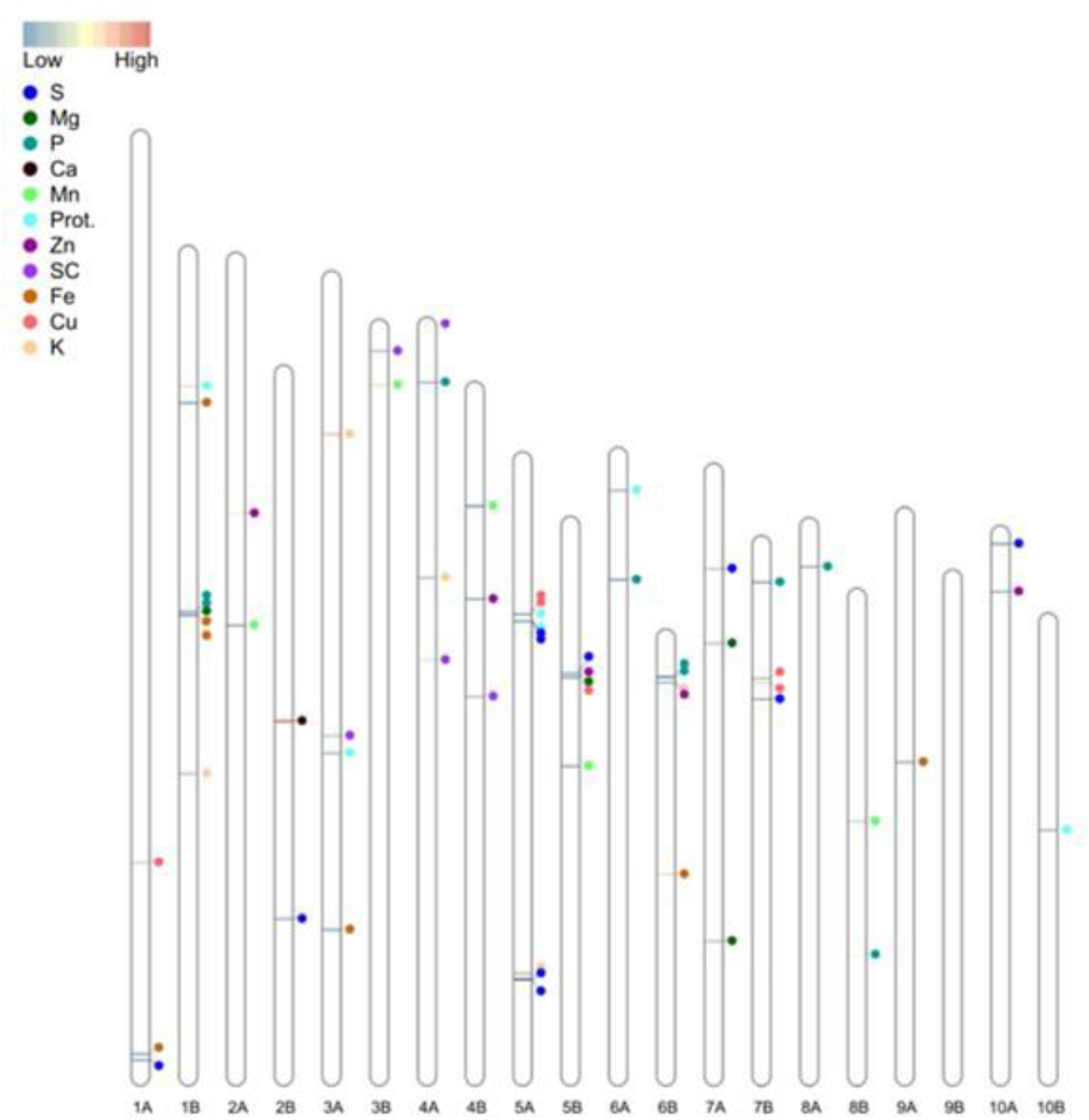
Distribution of 63 (significant and putative) marker–trait associations (MTAs) across the 20 tef chromosomes. Prot., protein; GC, seed color. The color of the horizontal bars on the chromosomes indicates the relative percentage of explained variation (PEV).

**Table 1.**
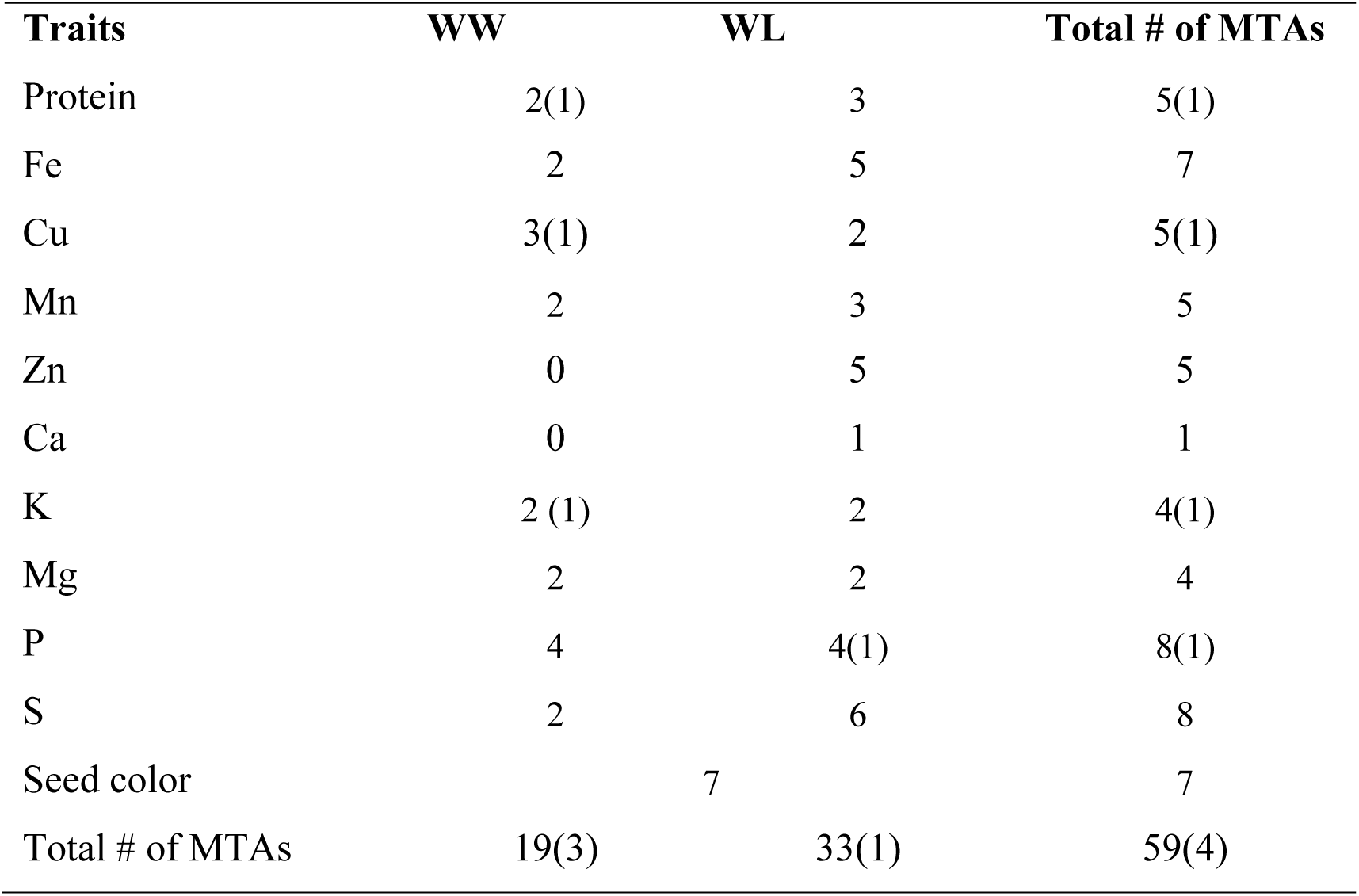
Summary of marker–trait associations (MTAs) detected for tef grain nutrient concentrations under well-watered (WW) and water-limited (WL) conditions in 2021. Number of significant MTAs meeting the Bonferroni-corrected threshold (*p* < 1.7E-06) is indicated, and number of putative MTAs, meeting a less stringent threshold (*p* < 1E-04) associated with the same SNPs is indicated in parentheses.

**Table 2.**
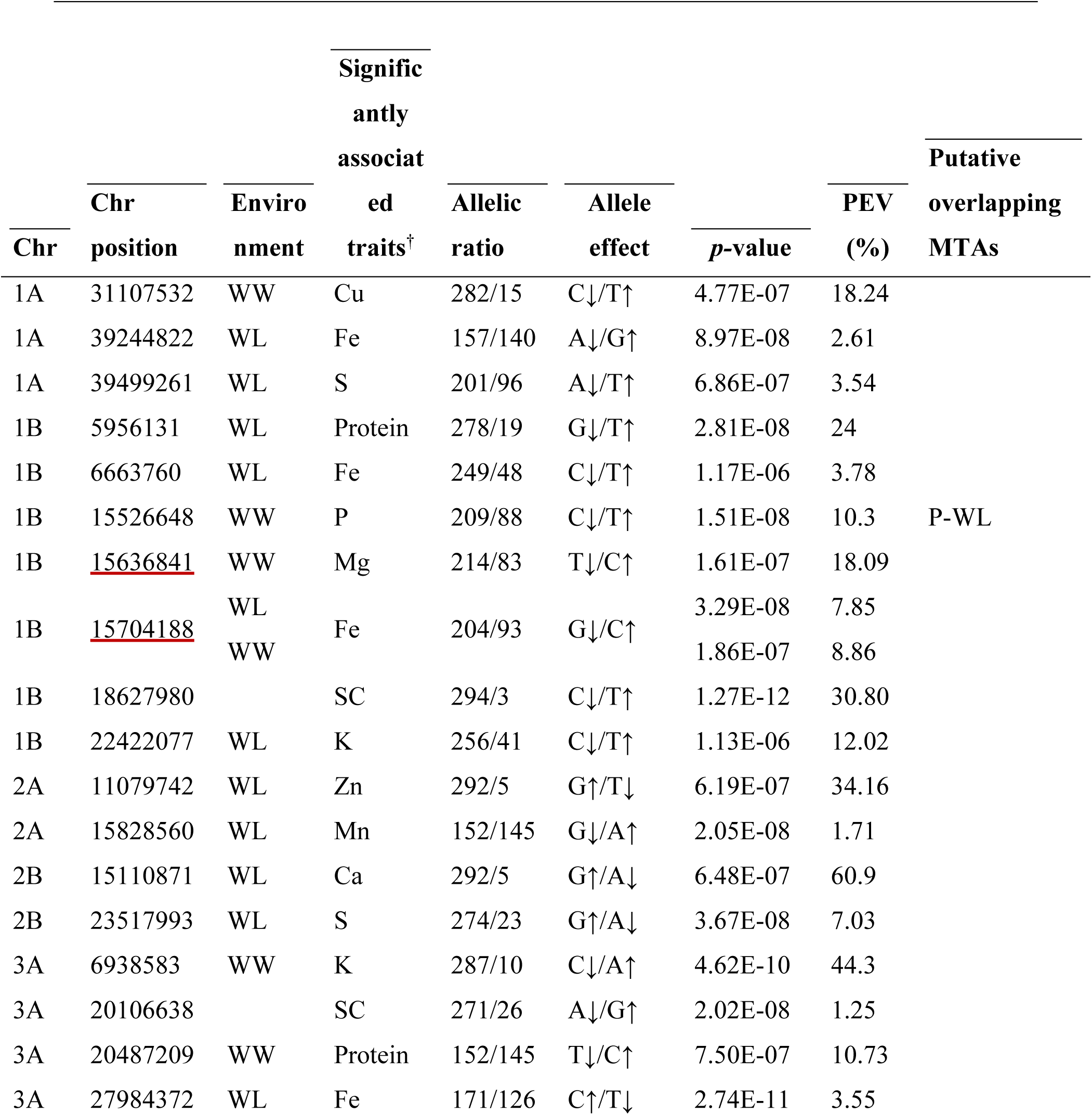
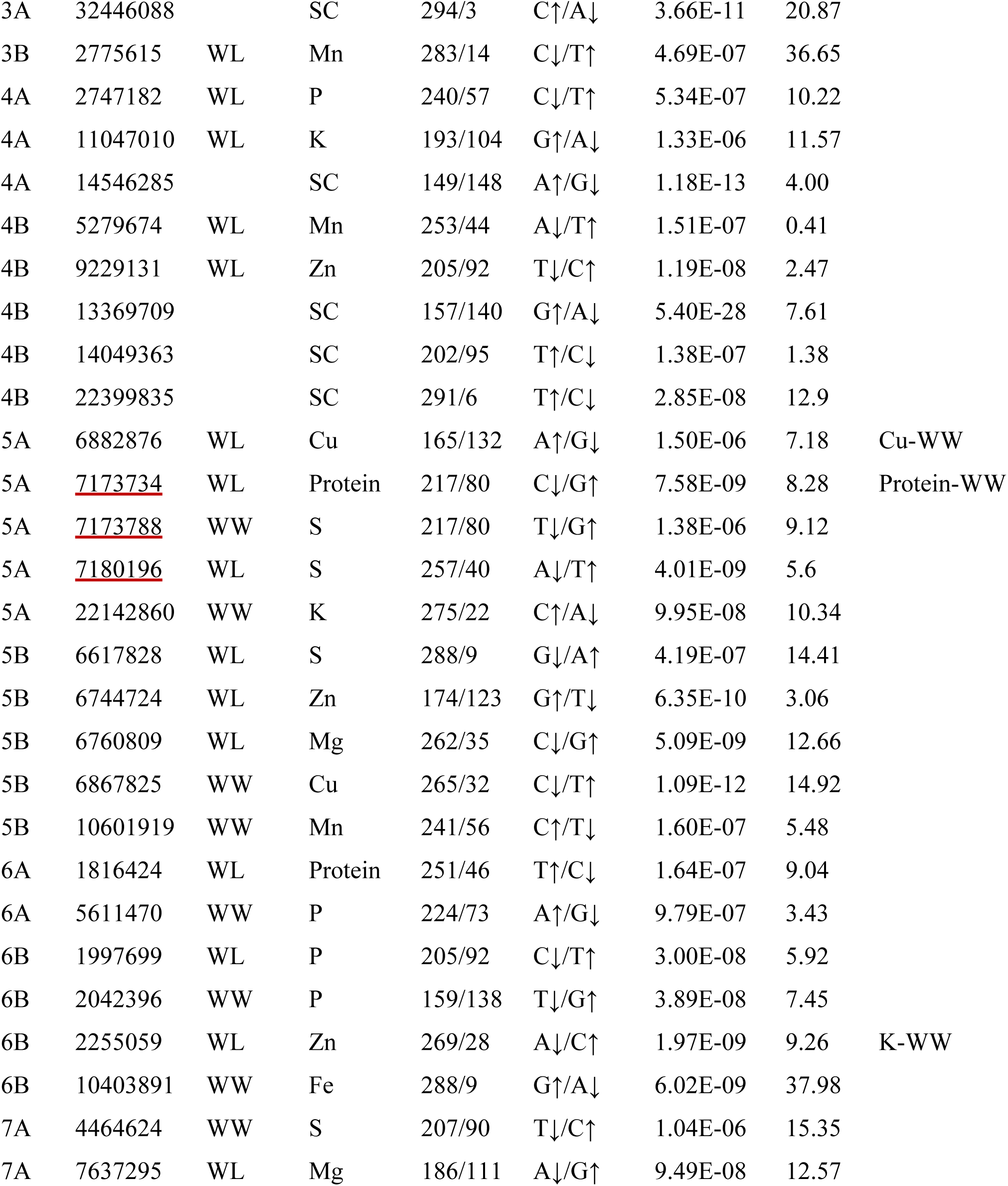
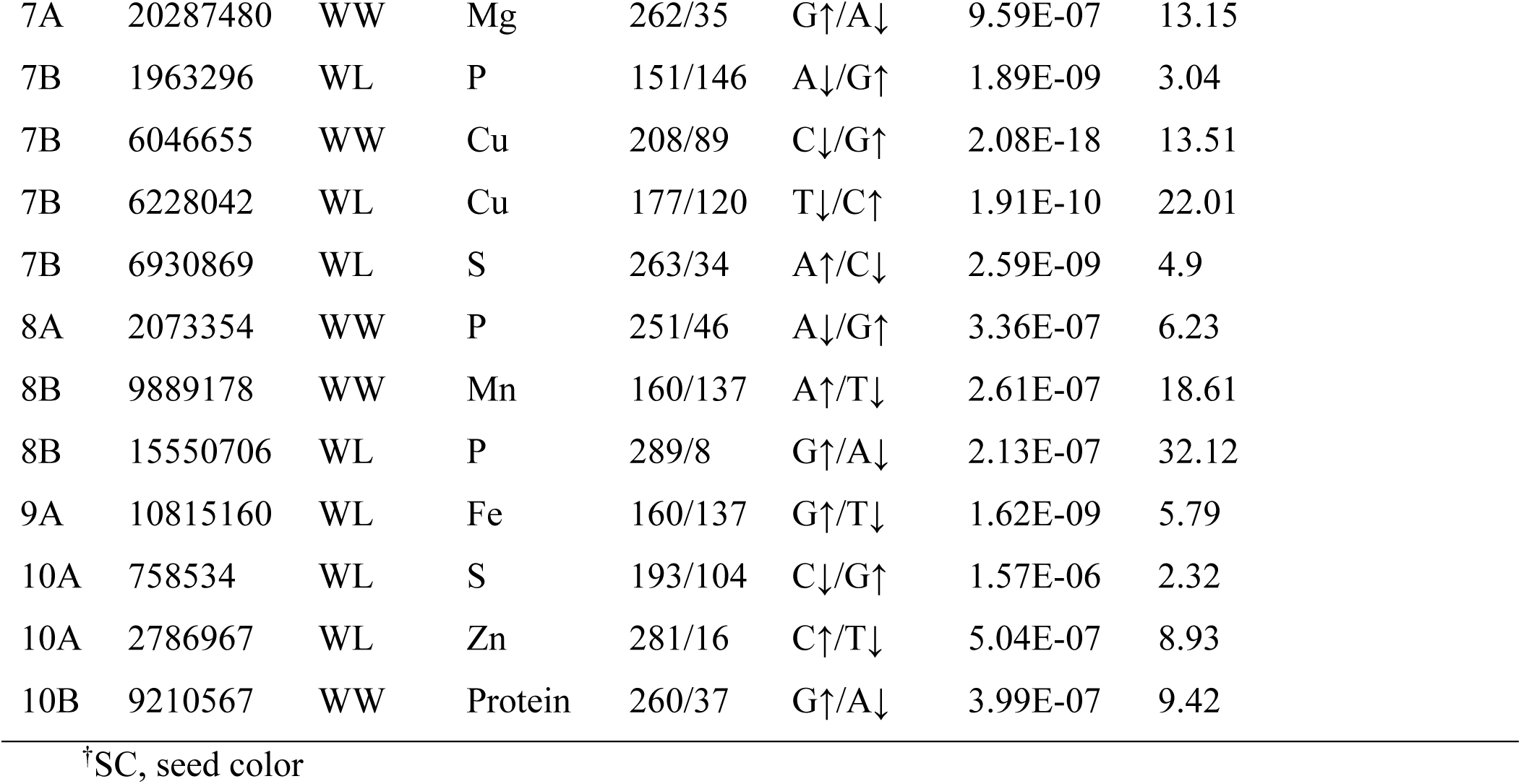
List of marker–trait associations (MTAs) detected for tef seed color and nutrient concentrations under well-watered (WW) and water-limited (WL) conditions in 2021. Significant MTAs, meeting the Bonferroni-corrected threshold (*p* < 1.7E-06), and putative MTAs meeting a less stringent threshold (*p* < 1E-04) and associated with the same SNPs. Details of MTAs include chromosome (Chr) number and position, environment, trait, allelic ratio, allele effect [increasing (↑) and decreasing (↓)], significance level (*p*-value) and percentage of explained variation (PEV). Closely linked MTAs, i.e., with physical distance < 97,242 bp and linkage r^2^ > 0.5 are underlined.

The detected (significand and putative) MTAs involved 58 SNPs, of which 53 were associated with a single trait under one environment, 4 were associated with the same trait under two environments, and 1 was associated with Zn and K (pleiotropic effect) (Table 2). LD analysis (Figure S4) detected two groups of closely linked of MTAs (underlined in Table 2). Among them, the 1B linkage group (two SNPs) was associated with Fe under both environments and Mg under the WW treatment, whereas the 5A linkage group (three SNPs) was associated with S and protein, both under the two environments.

Five significant (and one putative) MTAs were found for grain protein (Tables 1 and 2), with two of them involving the same SNP (5A_7173734) under both treatments (Table 2). The significant MTAs detected for protein explained 8.28–24% of the phenotypic variation, with the minor allele increasing protein concentration in three cases and the major allele increasing protein concentration in two cases.

A total of 22 significant (and one putative) MTAs were detected for grain micronutrients (Tables 1 and 2). Two SNPs (1B_15704188 and 5A_6882876) were each associated with the same mineral (Fe and Cu, respectively) under both treatments. The MTAs detected for micronutrients explained between 0.4% and 37.98% of the phenotypic variation, with the lowest and highest variation observed for Mn and Fe, respectively. In 13 cases, the minor alleles increased the concentration of grain micronutrients, whereas in 9 cases, the major alleles increased the concentration of micronutrients.

A total of 25 significant (and two putative) MTAs were detected for macronutrients (Tables 1 and 2). One SNP (1B_15526648) was associated with P under both treatments. The MTAs associated with macronutrients explained 2.3% to 60.9% of the phenotypic variation, with the lowest and highest variation observed for S and Ca, respectively. In most cases, the minor alleles increased the concentration of macronutrients (Table 2).

Seven significant MTAs were detected for seed color (Tables 1 and 2). The significant MTAs explained 11% to 29.36% of the phenotypic variation for this trait. In most cases, the major alleles were associated with white seeds, whereas the minor alleles were associated with brown seeds (Table 2).

## 4 Discussion

Global human diet is highly reliant on cereal crops, changes in their yield or quality can affect the food and nutrition security of millions of people (Galani et al., 2022, IFPRI, 2024). Drought stress slows grain-filling rate and shortens filling duration, thereby decreasing crop yields and nutritional properties (Sehgal et al., 2018). The effects of drought on yield and quality may vary depending on the crop species, genotype, environment, growth stage and stress severity (El Sabagh et al., 2020; Stagnari et al., 2016). While the effects of drought stress on the productivity of various crops have been extensively studied, the impact on grain quality and its genomic basis have been less investigated (Chadalavada et al., 2022; Stagnari et al., 2016). Recent development of high-throughput tools has created opportunities to examine the effects of drought on grain nutrient concentrations (Chadalavada et al., 2022; El Sabagh et al., 2020; Kamal et al., 2023). Nevertheless, there are only a few published studies on the effects of environmental factors on tef nutritional properties (Abewa et al., 2019) and their underlying genomic loci (Ereful et al., 2022), usually based on a small number of genotypes. Hence, the current study seems to be the first to investigate tef grain quality in a large number of genotypes under contrasting environments (WW and WL), and to implement a GWAS to detect the associated genomic loci.

### 4.1 Genetic diversity in grain nutrient concentrations

Grain mineral concentrations are the outcome of various physiological processes, including root uptake, translocation, redistribution within the plant tissues, remobilization and accumulation in the grain (Cakmak & Kutman, 2018; Marcos-Barbero et al., 2021). In the current study, grain nutrient concentrations exhibited wide variation among the tested genotypes across the two irrigation treatments (WW and WL) (Figure 1), thus reflecting the genetic diversity in tef germplasm. In previous studies, grain nutrient concentrations varied significantly across tef accessions in the field (Ereful et al., 2022) and greenhouse (Ligaba-Osena et al., 2021). The ranges of protein, micronutrients and macronutrients recorded in our study are mostly in agreement with previous reports on grain nutritional properties of tef grown under common field conditions in Ethiopia (Abewa et al., 2019; Daba, 2017), under irrigated field conditions in Israel (Tietel et al., 2020) and in greenhouse in Washington, USA (Ligaba-Osena et al., 2021). A clear deviation from this general agreement was found for Fe concentration, which in previous field studies (Abewa et al., 2019; Daba, 2017; Tietel et al., 2020), exhibited up to ∼20-fold the concentration found in our study, most probably due to soil contamination in the grain samples (Baye et al., 2014; Stangoulis & Sison, 2008). Nevertheless, our results confirm the superiority of tef grain mineral contents, including Fe, in comparison with published results for other cereal crops (Abebe et al., 2007; Baye et al., 2014; Dame, 2020; Ligaba-Osena et al., 2021).

Cereals crop grains have various colors, and contain substantial levels of pigments associated with grain nutrient content (Francavilla & Joye, 2020). Significant variations in nutrient concentrations have been observed between colored grains in maize (Martínez-Martínez et al., 2019) and wheat (Li et al., 2023). In the current study, differently colored tef grains differed significantly in all nutrient concentrations except Fe (Figure S1, Table S1), suggesting either a physiological or genetic association between grain color and nutrient uptake and/or accumulation. The variation in most nutrients’ concentrations between grain colors ranged from 0–7%, with the exception of Mn which exhibited 18% higher concentration in brown compared to white grains. Our results also indicated significantly higher Ca, Mg and P concentrations in the brown seeds, whereas protein, Cu, Zn, K and S were more plentiful in white-seeded genotypes. In previous studies, brown tef grain was found superior to white grain with respect to most minerals (Abebe et al., 2007; Dame, 2020; Tietel et al., 2020), while protein concentration was higher in white tef grain (Gebru et al., 2019).

Soil moisture represents a key physical factor affecting directly transport of nutrients to root surfaces through mass flow and diffusion in soils, root-nutrient contact and uptake and shoot transport (Brouder and Volenec, 2023; Coskun and White, 2023). In wheat, abiotic stress (drought and heat) negatively affects grain yield while increasing protein and mineral concentrations (Ben Mariem et al., 2021; Galani et al., 2022), however, in a wild emmer wheat collection no effect of drought was evident on protein, Fe or Zn content (Peleg et al., 2008). In the current study, water regimes had a significant effect on tef grain nutrient concentrations, with the exception of K (Figure 2, Table S1), which might be related to preferential deposition of K in the stem tissues of plants. Published reports show that up to 70 % of K in aboveground plant parts is transported and deposited in the stem tissue of various plants, such as wheat and rice (Xu et al., 2018; Zorb et al., 2014). Seasonal water application in our field experiment was reduced by ∼50% under WL compared to WW conditions, resulting in an average reduction of 42% in grain yield (Alemu et al., 2024). However, the effects of this severe drought treatment on grain quality attributes were relatively modest, usually ranging between 1 and 10% (Figure 2). Protein and most micronutrients exhibited higher concentrations under WL compared to WW conditions, providing partial compensation for the yield reduction, while most macronutrients exhibited the opposite trend. Among grain nutrients, Mn was the most influenced by water availability and exhibited a 25% lower concentration under the WL vs. WW treatment. It has been well-documented that root Mn uptake and shoot Mn accumulation are strongly affected by water regime. Most commonly, plant Mn concentrations show significant reductions with decreases in soil water status due to rapid oxidation of Mn to chemical forms unavailable for root uptake (Alejandro et al., 2020; Schmidt et al., 2016; Tao et al., 2007). It was previously reported that tef grain nutrient concentrations are influenced by the soil’s physicochemical properties (Abewa et al., 2019). However, we are not aware of any prior study on the effect of drought on tef grain quality.

In tef, GY was negatively associated with most grain nutrient concentrations under both treatments (Figure 3, Table S2), suggesting that higher yield under the WW treatment was counterbalanced by reduced nutrient concentration, as previously reported for wheat (Marcos-Barbero et al., 2021). In the current study, significant positive associations were observed in most cases under both treatments between grain protein and mineral concentrations, as well as among the various grain minerals. Notably, highly significant positive correlations were found between protein and Zn and protein and Fe (Table S2). Previously, it has been shown that Zn, Fe and protein are co-localized in the same grain fractions of wheat, leading to suggestion that grain protein acts as a sink for Zn and Fe (Cakmak et al., 2010). Similar correlations between grain mineral concentrations have been previously reported in tef (Ligaba-Osena et al., 2021) and wheat (Gomez-Becerra et al., 2010; Marcos-Barbero et al., 2021), suggesting synergic physiological mechanisms or common genetic control.

### 4.2 MTAs identified for grain nutrients

A total of 59 significant and 4 putative MTAs were identified for seed color and grain nutrients under WW and WL environments (Table 2, Figure S3). MTAs were distributed across 19 out of the 20 tef chromosomes, the exception being Chr 9B (Figure 4). Similar to our previous tef GWAS on productivity and drought-adaptive traits (Alemu et al., 2024), a greater number of significant MTAs were detected under the WL vs. WW treatment (Table 2). This suggests that genes induced by drought and possibly involved in drought-adaptive responses (Alemu et al., 2024) influence nutrient uptake and accumulation in the grains. GWAS has been used in various crops to identify loci underlying grain quality (Chandra et al., 2024; Kimani et al., 2020; Peleg et al., 2009; Puranik et al., 2020). However, reports on genomic dissection of nutritional quality in orphan crops are rather limited, and we are not aware of any previous genomic dissection of tef grain nutritional quality under contrasting water regimes.

### 4.3 Genomic loci highlight multiple-environment and pleiotropic effects

Five genomic regions reflected multiple-environment associations, including four cases of the same SNP (5A_7173734, 1B_15704188, 5A_6882876 and 1B_15526648) and one case of two closely linked SNPs (5A_7173788–7180196) associated with the same trait (protein, Fe, Cu, P and S, respectively) across our two environments. Similarly, multiple-environment associations have been identified in tef for productivity and drought-adaptive traits across irrigation regimes (Alemu et al., 2024), as well as for productivity and phenology across various locations in Ethiopia (Woldeyohannes et al., 2022). These multiple-environment associations reflect the effect of highly reliable constitutive genes underlying various grain quality traits, which are very likely to prove valuable for marker-assisted selection (McLeod et al., 2023) to improve tef adaptation to diverse environments.

Three genomic regions reflected pleiotropic effects, including one case on a single locus (6B_2255059) associated with both Zn (under WL) and K (under WW) and two groups of closely linked SNPs, 1B_15636841–15704188 and 5A_7173734–7180196, associated with Mg + Fe and protein + S, respectively. Whereas pleiotropism between Zn and K was supported by significant phenotypic association only under WL conditions (Figure 3, Table S2), the other two pleiotropic effects were supported by highly significant correlations across both environments. Similarly, pleiotropic loci were identified in TDP-300 in our previous study (Alemu et al., 2024). These pleiotropic relationships and significant associations suggest the existence of genetic correlations between traits (Alemu et al., 2024; Chebib & Guillaume, 2021; Mathew et al., 2019). Pleiotropy can indicate a shared etiology between traits (Lee et al., 2021; Von Berg et al., 2022) due to common genetic factors, such as the same gene(s) controlling both traits or closely linked genes (Chebib & Guillaume, 2021; Fang et al., 2017).

## 5 Concluding Remarks

Under the current climate change scenario, drought and heat stress are becoming the most severe constraints to crop production and quality, thus threating worldwide food and nutrition security. Therefore, a global focus on stress-resilient, high-yielding, nutritious crops is critical. Underutilized crops, such as tef, present an outstanding opportunity to increase crop diversity, promote climate resilience and boost nutritional quality. However, genomic and physiological dissection of tef under abiotic stress is still limited. In our previous study on tef, we focused on the genetic diversity and a genomic dissection of productivity and drought-adaptive traits (Alemu et al 2024a), as well as on the whole plant’s physiological responses to drought (Alemu et al. 2024b). These are supplemented by the current work on nutrient concentrations in tef grains under contrasting water regimes.

Grain nutrient concentration depends on genotype, environment and management, all contributing to the particularly wide variation between genotypes, but also between grain colors and irrigation regimes. Both seed color and irrigation treatment exhibited an inconsistent effect (either increasing or decreasing) on grain nutrients, generally up to 10% in magnitude. As discussed above, it is interesting to note that Mn exhibited the greatest responses to both water regime and seed color.

Genomic loci that reflect multiple-environment and pleiotropic associations provide new insights into the complex genetic basis of grain nutrients and the potential for breeding novel tef cultivars. The pleiotropism and phenotypic associations between grain nutrients may facilitate simultaneous selection and improvement of tef nutritional properties.

Our previous (Alemu et al., 2024) and current genomic dissection studies shed new light on tef responses to stress and may contribute to the main targets in tef breeding, as well as to further studies and a deeper understanding of tef genomics and physiology. Overall, tef productivity, drought-responsive traits and grain nutrient qualities are highly influenced by the environment and farming system. Although our studies were conducted in an irrigated Mediterranean environment, the genomic results may contribute to the development of drought-resistant, high-yielding and nutritious tef varieties, for Ethiopia and elsewhere.

## Acknowledgments

This study was supported by The Israel Innovation Authority, Challenge Program [grant no. 73546]. We gratefully acknowledge the Heinrich Bonnenberg Scholarship awarded to M.D.A., and the Robert H. Smith Foundation for doctoral fellowships awarded to M.D.A. and S.B.-Z. The authors wish to acknowledge the Israel Plant Gene Bank for providing the tef germplasm collection. We thank the Kvutzat Shiller farm for providing the platform for the field experiment. Y.S. is the incumbent of the Haim Gvati Chair in Agriculture.

## Author Contributions

MDA: Conceptualization, Data curation, Formal analysis, Investigation, Methodology, Writing – original draft, Writing – review & editing. SB-Z: Investigation, Writing – review & editing. VB: Investigation, Writing – review & editing. YT: Investigation, Methodology, Writing – review & editing. IC: Investigation, Methodology, Writing – review & editing. YS: Conceptualization, Formal analysis, Funding acquisition, Investigation, Methodology, Project administration, Supervision, Validation, Writing – review & editing.

## Conflict of Interest

The authors declare no conflict of interest.

## Germplasm and Data Availability Statement

Seeds of the tef diversity panel (TDP-300) are available upon reasonable request from the corresponding author. Data supporting the findings will be available: for the raw DNA sequencing reads, Short Read Archive (SRA) (https://www.ncbi.nlm.nih.gov/sra/PRJNA1063642, BioProject accession number PRJNA1063642); for the SNP and phenotypic data, Zenodo (https://doi.org/10.5281/zenodo.10508959).

## **7** Supplementary information

**Figure S1.**
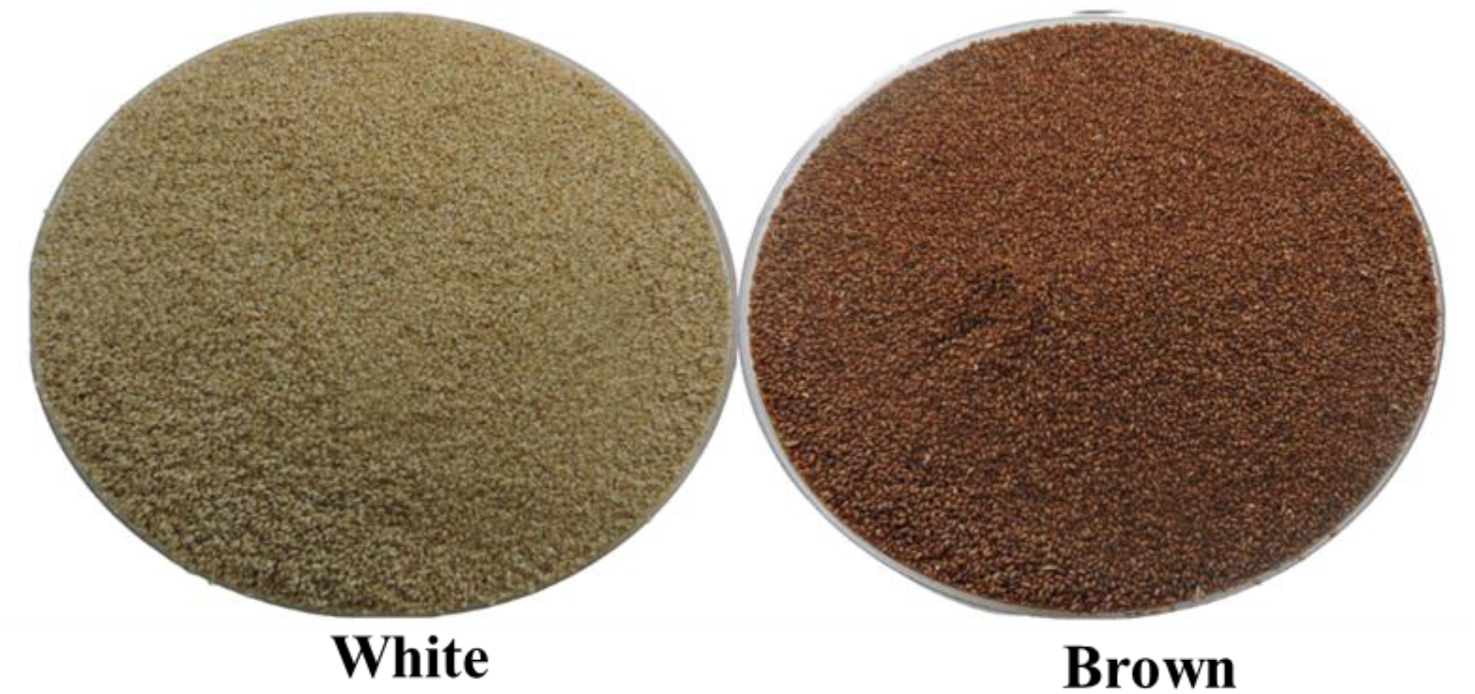
Tef diversity panel (TDP-300) grain colors visually scored into two broad categories: white and brown.

**Figure S2.**
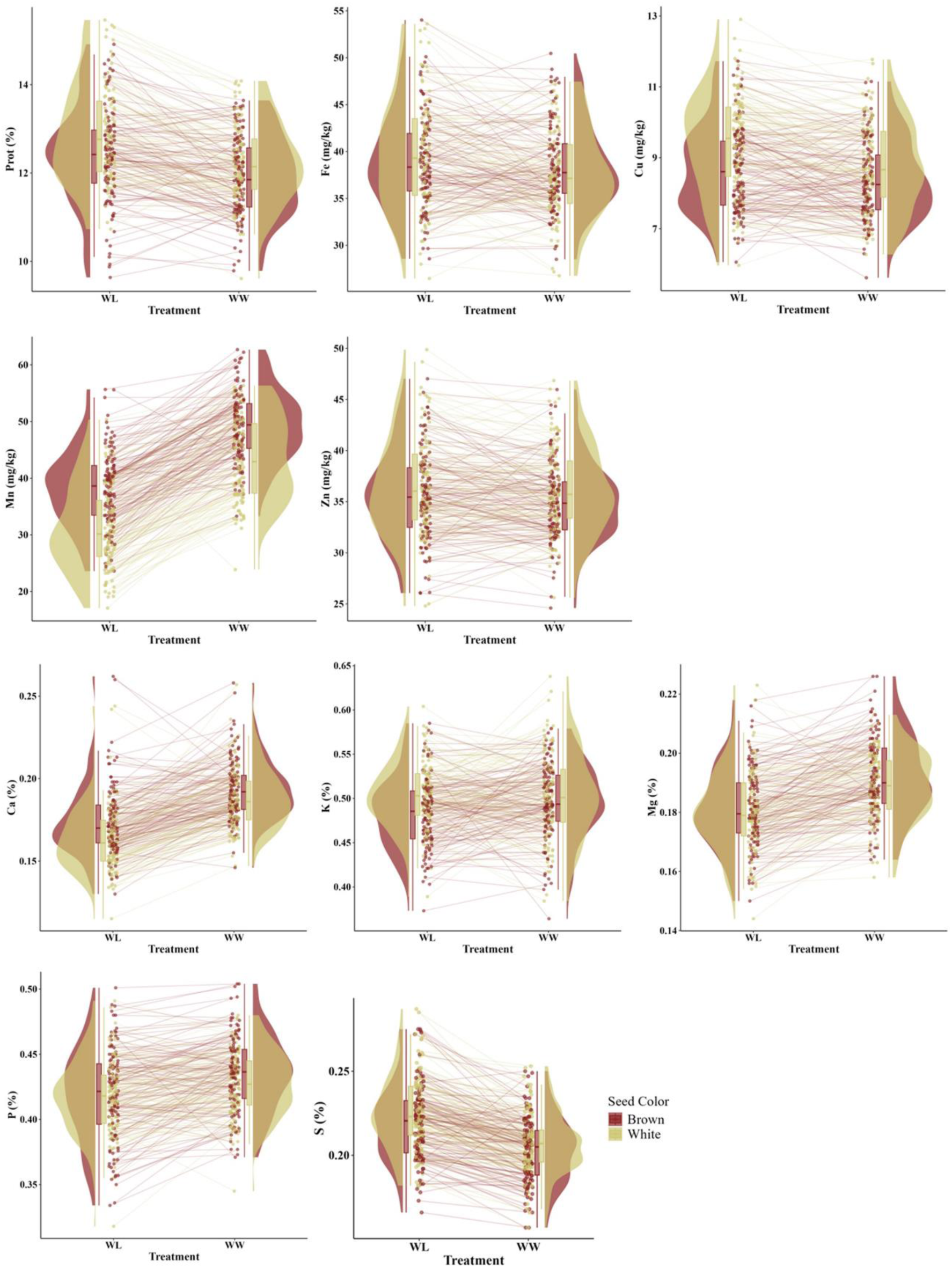
Raincloud plots depicting the effect of genotype (brown and white seed colors)-by-environment (well-watered, WW; water-limited, WL) interaction on the concentrations of tef grain nutrients in 2021. Prot, protein.

**Figure S3.**
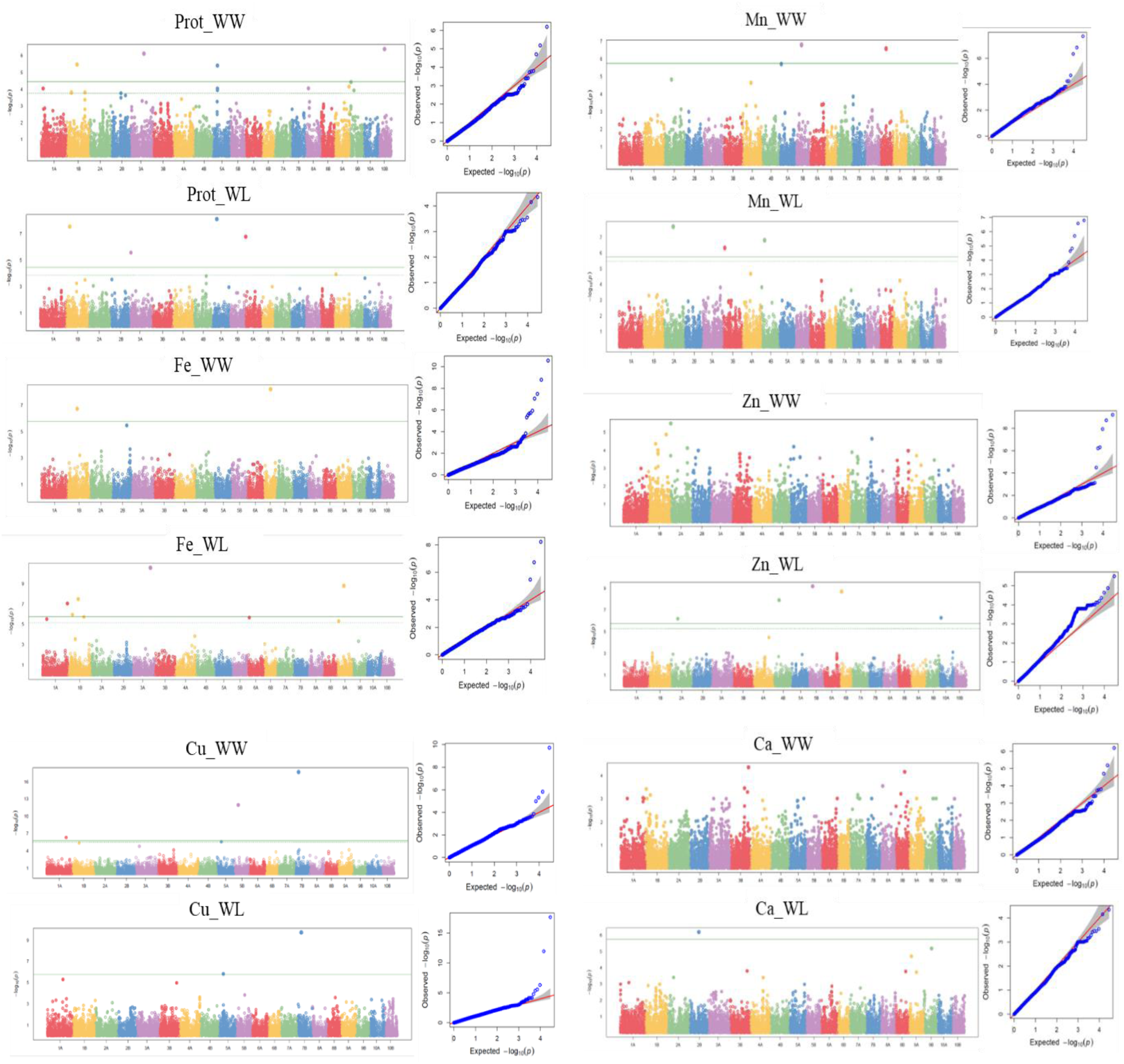
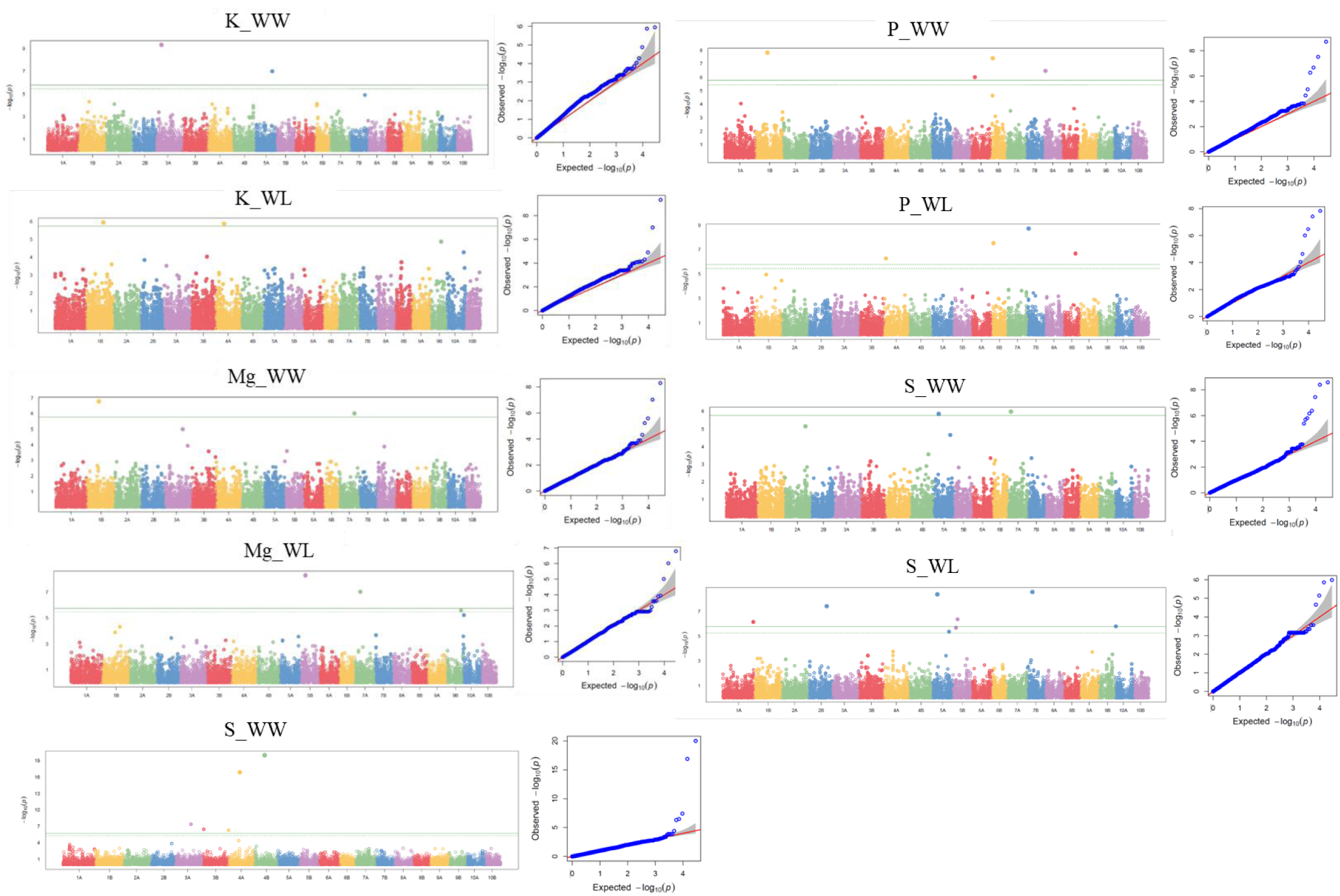
Genome-wide association study (GWAS) in tef. Manhattan plot showing 20 chromosomes with significant single-nucleotide polymorphisms (SNPs) associated with grain nutritional content and quantile-quantile (Q-Q) plot under well-watered (WW) and water-limited (WL) treatments in 2021. Prot, protein. Bonferroni threshold, *p* < 1.7E-06 (horizontal solid green line); less stringent threshold, *p* < 1E-04 (dashed green line). X-axis represents the 20 tef chromosomes.

**Figure S4.**
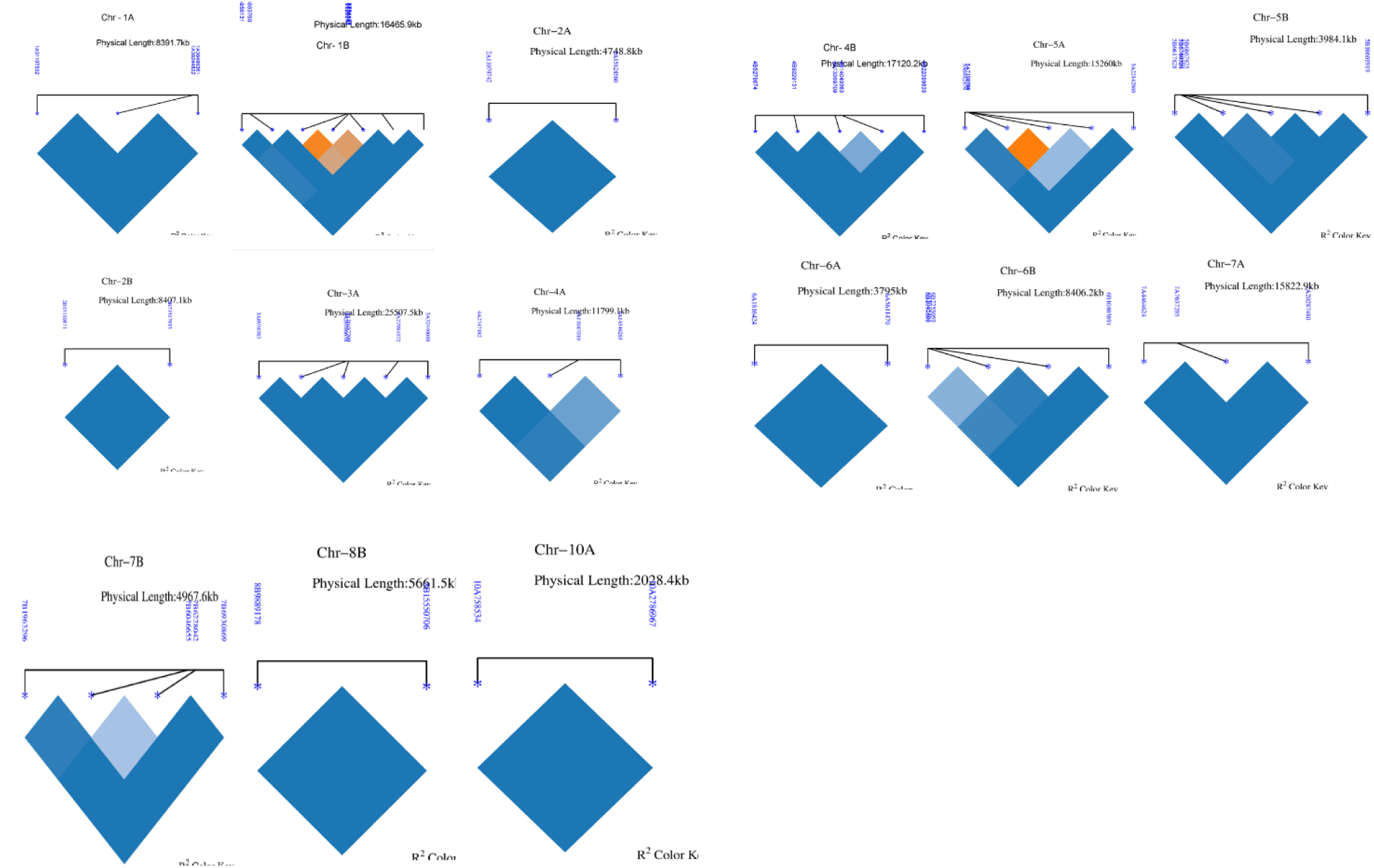
Graphical display of pairwise linkage disequilibrium (LD) heat map of 59 significant single-nucleotide polymorphisms (SNPs) on 15 out of 20 chromosomes (Chr) (excluding Chr 3B, 8A, 9A & B and 10B which exhibited single significant SNP).

## Notes

### Competing Interest Statement

The authors have declared no competing interest.

## 6 References

Abebe, Y., Bogale, A., Hambidge, K. M., Stoecker, B. J., Bailey, K., & Gibson, R. S. (2007). Phytate, zinc, iron and calcium content of selected raw and prepared foods consumed in rural Sidama, Southern Ethiopia, and implications for bioavailability. Journal of Food Composition and Analysis, 20(3–4), 161–168. 10.1016/j.jfca.2006.09.003

Abewa, A., Adgo, E., Yitaferu, B., Alemayehu, G., Assefa, K., Solomon, J. K. Q., & Payne, W. (2019). Teff Grain Physical and Chemical Quality Responses to Soil Physicochemical Properties and the Environment. Agronomy, 9(6), 283. 10.3390/agronomy9060283

Alejandro, S., Höller, S., Meier, B., & Peiter, E. (2020). Manganese in Plants: From Acquisition to Subcellular Allocation. Frontiers in Plant Science, 11, 300. 10.3389/fpls.2020.00300

Alemu, M. D., Ben-Zeev, S., Hellwig, T., Barak, V., Shoshani, G., Chen, A., Razzon, S., Herrmann, I., Vorobyova, A., Hübner, S., & Saranga, Y. (2024). Genomic dissection of productivity, lodging, and morpho-physiological traits in *ERAGROSTIS TEF* under contrasting water availabilities. PLANTS, PEOPLE, PLANET, ppp3.10505. 10.1002/ppp3.10505

Allaby, R. (2021). First come, first served for ancient crops. Nature Plants, 7(5), 542–543. 10.1038/s41477-021-00926-w

Allen, M., Poggiali, D., Whitaker, K., Marshall, T. R., Van Langen, J., & Kievit, R. A. (2021). Raincloud plots: A multi-platform tool for robust data visualization. Wellcome Open Research, 4, 63. 10.12688/wellcomeopenres.15191.2

Asefa, B. G., Tsige, F., Mehdi, M., Kore, T., & Lakew, A. (2023). Rapid classification of tef [Eragrostis tef (Zucc.) Trotter] grain varieties using digital images in combination with multivariate technique. Smart Agricultural Technology, 3, 100097. 10.1016/j.atech.2022.100097

Assefa, K., Chanyalew, S., & Tadele, Z. (2017). Tef, *Eragrostis tef* (Zucc.) Trotter. In J. V. Patil (Ed.), Millets and Sorghum (1st ed., pp. 226–266). Wiley. 10.1002/9781119130765.ch9

Assefa, K., Yu, J.-K., Zeid, M., Belay, G., Tefera, H., & Sorrells, M. E. (2011). Breeding tef [*Eragrostis tef* (Zucc.) trotter]: Conventional and molecular approaches. Plant Breeding, 130(1), 1–9. 10.1111/j.1439-0523.2010.01782.x

Atwell, S., Huang, Y. S., Vilhjálmsson, B. J., Willems, G., Horton, M., Li, Y., Meng, D., Platt, A., Tarone, A. M., Hu, T. T., Jiang, R., Muliyati, N. W., Zhang, X., Amer, M. A., Baxter, I., Brachi, B., Chory, J., Dean, C., Debieu, M., … Nordborg, M. (2010). Genome-wide association study of 107 phenotypes in Arabidopsis thaliana inbred lines. Nature, 465(7298), 627–631. 10.1038/nature08800

Baye, K., Mouquet-Rivier, C., Icard-Vernière, C., Picq, C., & Guyot, J. (2014). Changes in mineral absorption inhibitors consequent to fermentation of E thiopian *injera*: Implications for predicted iron bioavailability and bioaccessibility. International Journal of Food Science & Technology, 49(1), 174–180. 10.1111/ijfs.12295

Bekkering, C. S., & Tian, L. (2019). Thinking Outside of the Cereal Box: Breeding Underutilized (Pseudo)Cereals for Improved Human Nutrition. Frontiers in Genetics, 10, 1289. 10.3389/fgene.2019.01289

Ben Mariem, S., Soba, D., Zhou, B., Loladze, I., Morales, F., & Aranjuelo, I. (2021). Climate Change, Crop Yields, and Grain Quality of C3 Cereals: A Meta-Analysis of [CO2], Temperature, and Drought Effects. Plants, 10(6), 1052. 10.3390/plants10061052

Bonferroni, C. (1936). Teoria statistica delle classi e calcolo delle probabilita. Pubblicazioni Del R Istituto Superiore Di Scienze Economiche e Commericiali Di Firenze, 8, 3–62.

Brouder SM, Valenec JJ (2023) Nutrition of plants in a changing climate. In: Rengel Z, Cakmak I, White PJ (eds) Marschner’s Mineral Nutrition of Plants, 4th edn. Academic Press Elsevier Ltd, pp 723–750. 10.1016/B978-0-12-819773-8.00023X

Cakmak, I., & Kutman, U. B. (2018). Agronomic biofortification of cereals with zinc: A review. European Journal of Soil Science, 69(1), 172–180. 10.1111/ejss.12437

Cakmak, I., Pfeiffer, W. H., & McClafferty, B. (2010). REVIEW: Biofortification of Durum Wheat with Zinc and Iron. Cereal Chemistry, 87(1), 10–20. 10.1094/CCHEM-87-1-0010

Chadalavada, K., Guna, K., Kumari, B. R., & Kumar, T. S. (2022). Drought stress in sorghum: Impact on grain quality. In Climate Change and Crop Stress (pp. 113–134). Elsevier.

Chandra, A. K., Pandey, D., Sood, S., Joshi, D. C., Tiwari, A., Sharma, D., Gururani, K., & Kumar, A. (2024). Uncovering the genomic regions underlying grain iron and zinc content using genome-wide association mapping in finger millet. *3* Biotech, 14(2), 47. 10.1007/s13205-023-03889-1

Chanyalew, S., Ferede, S., Damte, T., Fikre, T., Genet, Y., Kebede, W., Tolossa, K., Tadele, Z., & Assefa, K. (2019). Significance and prospects of an orphan crop tef. Planta, 250(3), 753–767. 10.1007/s00425-019-03209-z

Chapman, M. A., He, Y., & Zhou, M. (2022). Beyond a reference genome: Pangenomes and population genomics of underutilized and orphan crops for future food and nutrition security. New Phytologist, 234(5), 1583–1597. 10.1111/nph.18021

Chebib, J., & Guillaume, F. (2021). Pleiotropy or linkage? Their relative contributions to the genetic correlation of quantitative traits and detection by multitrait GWA studies. Genetics, 219(4), iyab159. 10.1093/genetics/iyab159

Cheng, A. (2018). Review: Shaping a sustainable food future by rediscovering long-forgotten ancient grains. Plant Science, 269, 136–142. 10.1016/j.plantsci.2018.01.018

Coskun, D., White, P.J., 2023. Ion-uptake mechanisms of individual cells and roots: short-distance transport. In: Rengel Z, Cakmak I, White PJ (eds) Marschner’s Mineral Nutrition of Plants, 4th edn. Academic Press Elsevier Ltd, pp 723–750. 10.1016/B978-0-12-819773-8.00023-X

Daba, T. (2017). Nutritional and soio-cultural values of teff (Eragrostis tef) in Ethiopia. International Journal of Food Science and Nutrition, 2(3), 50–57.

Dame, Z. T. (2020). Analysis of major and trace elements in teff (Eragrostis tef). Journal of King Saud University - Science, 32(1), 145–148. 10.1016/j.jksus.2018.03.020

D’Andrea, A. C. (2008). T’ef (Eragrostis tef) in Ancient Agricultural Systems of Highland Ethiopia. Economic Botany, 62(4), 547–566. 10.1007/s12231-008-9053-4

Dietz, K.-J., Zörb, C., & Geilfus, C.-M. (2021). Drought and crop yield. Plant Biology, 23(6), 881–893. 10.1111/plb.13304

El Sabagh, A., Hossain, A., Barutçular, C., Islam, M. S., Ahmad, Z., Wasaya, A., Meena, R. S., Fahad, S., Oksana, S., Hafez, Y. M., Najeeb, U., Çiğ, F., Konuşkan, Ö., & Hasanuzzaman, M. (2020). Adverse Effect of Drought on Quality of Major Cereal Crops: Implications and Their Possible Mitigation Strategies. In M. Hasanuzzaman (Ed.), Agronomic Crops (pp. 635–658). Springer Singapore. 10.1007/978-981-15-0025-1_31

Ereful, N. C., Jones, H., Fradgley, N., Boyd, L., Cherie, H. A., & Milner, M. J. (2022). Nutritional and genetic variation in a core set of Ethiopian Tef (Eragrostis tef) varieties. BMC Plant Biology, 22(1), 220. 10.1186/s12870-022-03595-9

Fang, C., Ma, Y., Wu, S., Liu, Z., Wang, Z., Yang, R., Hu, G., Zhou, Z., Yu, H., Zhang, M., Pan, Y., Zhou, G., Ren, H., Du, W., Yan, H., Wang, Y., Han, D., Shen, Y., Liu, S., … Tian, Z. (2017). Genome-wide association studies dissect the genetic networks underlying agronomical traits in soybean. Genome Biology, 18(1), 161. 10.1186/s13059-017-1289-9

FAO. (2020). The State of Food Security and Nutrition in the World 2020. FAO, IFAD, UNICEF, WFP and WHO. 10.4060/ca9692en

FAO. (2023). The State of Food Security and Nutrition in the World 2023. FAO; IFAD; UNICEF; WFP; WHO; 10.4060/cc3017en

Francavilla, A., & Joye, I. J. (2020). Anthocyanins in Whole Grain Cereals and Their Potential Effect on Health. Nutrients, 12(10), 2922. 10.3390/nu12102922

Galani, Y. J. H., Hansen, E. M. Ø., Droutsas, I., Holmes, M., Challinor, A. J., Mikkelsen, T. N., & Orfila, C. (2022). Effects of combined abiotic stresses on nutrient content of European wheat and implications for nutritional security under climate change. Scientific Reports, 12(1), 5700. 10.1038/s41598-022-09538-6

Gebru, Y. A., Hyun-II, J., Young-Soo, K., Myung-Kon, K., & Kwang-Pyo, K. (2019). Variations in Amino Acid and Protein Profiles in White versus Brown Teff (Eragrostis Tef) Seeds, and Effect of Extraction Methods on Protein Yields. Foods, 8(6), 202. 10.3390/foods8060202

Gomez-Becerra, H. F., Erdem, H., Yazici, A., Tutus, Y., Torun, B., Ozturk, L., & Cakmak, I. (2010). Grain concentrations of protein and mineral nutrients in a large collection of spelt wheat grown under different environments. Journal of Cereal Science, 52(3), 342–349. 10.1016/j.jcs.2010.05.003

Hager, A.-S., Wolter, A., Jacob, F., Zannini, E., & Arendt, E. K. (2012). Nutritional properties and ultra-structure of commercial gluten free flours from different botanical sources compared to wheat flours. Journal of Cereal Science, 56(2), 239–247. 10.1016/j.jcs.2012.06.005

Huang, M., Liu, X., Zhou, Y., Summers, R. M., & Zhang, Z. (2019). BLINK: A package for the next level of genome-wide association studies with both individuals and markers in the millions. GigaScience, 8(2). 10.1093/gigascience/giy154

Huang, X., & Han, B. (2014). Natural Variations and Genome-Wide Association Studies in Crop Plants. Annual Review of Plant Biology, 65(1), 531–551. 10.1146/annurev-arplant-050213-035715

Hunter, D., Borelli, T., Beltrame, D. M. O., Oliveira, C. N. S., Coradin, L., Wasike, V. W., Wasilwa, L., Mwai, J., Manjella, A., Samarasinghe, G. W. L., Madhujith, T., Nadeeshani, H. V. H., Tan, A., Ay, S. T., Güzelsoy, N., Lauridsen, N., Gee, E., & Tartanac, F. (2019). The potential of neglected and underutilized species for improving diets and nutrition. Planta, 250(3), 709–729. 10.1007/s00425-01903169-4

International Food Policy Research Institute (IFPRI). 2024. Global food policy report 2024: Food systems for healthy diets and nutrition. Washington, DC: International Food Policy Research Institute. https://hdl.handle.net/10568/141760

Islam, A. S. M. F., Mustahsan, W., Tabien, R., Awika, J. M., Septiningsih, E. M., & Thomson, M. J. (2022). Identifying the Genetic Basis of Mineral Elements in Rice Grain Using Genome-Wide Association Mapping. Genes, 13(12), 2330. 10.3390/genes13122330

Jaiswal, V., Bandyopadhyay, T., Gahlaut, V., Gupta, S., Dhaka, A., Ramchiary, N., & Prasad, M. (2019). Genome-wide association study (GWAS) delineates genomic loci for ten nutritional elements in foxtail millet (Setaria italica L.). Journal of Cereal Science, 85, 48–55. 10.1016/j.jcs.2018.11.006

Jifar, H., Assefa, K., & Tadele, Z. (2015). Grain yield variation and association of major traits in brown-seeded genotypes of tef [Eragrostis tef (Zucc.)Trotter]. Agriculture & Food Security, 4(1), 7. 10.1186/s40066-015-0027-3

Kamal, N. M., Gorafi, Y. S. A., Tomemori, H., Kim, J.-S., Elhadi, G. M. I., & Tsujimoto, H. (2023). Genetic variation for grain nutritional profile and yield potential in sorghum and the possibility of selection for drought tolerance under irrigated conditions. BMC Genomics, 24(1), 515. 10.1186/s12864-023-09613-w

Kimani, W., Zhang, L.-M., Wu, X.-Y., Hao, H.-Q., & Jing, H.-C. (2020). Genome-wide association study reveals that different pathways contribute to grain quality variation in sorghum (Sorghum bicolor). BMC Genomics, 21(1), 112. 10.1186/s12864-020-6538-8

Lee, C. H., Shi, H., Pasaniuc, B., Eskin, E., & Han, B. (2021). PLEIO: A method to map and interpret pleiotropic loci with GWAS summary statistics. The American Journal of Human Genetics, 108(1), 36–48. 10.1016/j.ajhg.2020.11.017

Li, L., Zhang, H., Liu, J., Huang, T., Zhang, X., Xie, H., Guo, Y., Wang, Q., Zhang, P., & Qin, P. (2023). Grain color formation and analysis of correlated genes by metabolome and transcriptome in different wheat lines at maturity. Frontiers in Nutrition, 10, 1112497. 10.3389/fnut.2023.1112497

Ligaba-Osena, A., Mengistu, M., Beyene, G., Cushman, J., Glahn, R., & Piñeros, M. (2021). Grain mineral nutrient profiling and iron bioavailability of an ancient crop tef (Eragrostis tef). Australian Journal of Crop Science, 15(*10*):2021, 1314–1324. 10.21475/ajcs.21.15. 10.p3264

Liu, J., Huang, L., Li, T., Liu, Y., Yan, Z., Tang, G., Zheng, Y., Liu, D., & Wu, B. (2021). Genome-Wide Association Study for Grain Micronutrient Concentrations in Wheat Advanced Lines Derived From Wild Emmer. Frontiers in Plant Science, 12, 651283. 10.3389/fpls.2021.651283

Mabhaudhi, T., Chimonyo, V. G. P., Hlahla, S., Massawe, F., Mayes, S., Nhamo, L., & Modi, A. T. (2019). Prospects of orphan crops in climate change. Planta, 250(3), 695–708. 10.1007/s00425-019-03129-y

Marcos-Barbero, E. L., Pérez, P., Martínez-Carrasco, R., Arellano, J. B., & Morcuende, R. (2021). Genotypic Variability on Grain Yield and Grain Nutritional Quality Characteristics of Wheat Grown under Elevated CO2 and High Temperature. Plants, 10(6), 1043. 10.3390/plants10061043

Martínez-Martínez, R., Chávez-Servia, J. L., Vera-Guzmán, A. M., Aquino-Bolaños, E. N., Carrillo-Rodríguez, J. C., & Pérez-Herrera, A. (2019). Phenotypic variation in grain mineral compositions of pigmented maize conserved in indigenous communities of Mexico.

Mathew, I., Shimelis, H., Shayanowako, A. I. T., Laing, M., & Chaplot, V. (2019). Genome-wide association study of drought tolerance and biomass allocation in wheat. PLOS ONE, 14(12), e0225383. 10.1371/journal.pone.0225383

McLeod, L., Barchi, L., Tumino, G., Tripodi, P., Salinier, J., Gros, C., Boyaci, H. F., Ozalp, R., Borovsky, Y., Schafleitner, R., Barchenger, D., Finkers, R., Brouwer, M., Stein, N., Rabanus-Wallace, M. T., Giuliano, G., Voorrips, R., Paran, I., & Lefebvre, V. (2023). Multi-environment association study highlights candidate genes for robust agronomic quantitative trait loci in a novel worldwide *Capsicum* core collection. The Plant Journal, 116(5), 1508–1528. 10.1111/tpj.16425

Milla, R., & Osborne, C. P. (2021). Crop origins explain variation in global agricultural relevance. Nature Plants, 7(5), 598–607. 10.1038/s41477-021-00905-1

Nyiraguhirwa, S., Grana, Z., Ouabbou, H., Iraqi, D., Ibriz, M., Mamidi, S., & Udupa, S. M. (2022). A Genome-Wide Association Study Identifying Single-Nucleotide Polymorphisms for Iron and Zinc Biofortification in a Worldwide Barley Collection. Plants, 11(10), 1349. 10.3390/plants11101349

Peleg, Z., Cakmak, I., Ozturk, L., Yazici, A., Jun, Y., Budak, H., Korol, A. B., Fahima, T., & Saranga, Y. (2009). Quantitative trait loci conferring grain mineral nutrient concentrations in durum wheat × wild emmer wheat RIL population. Theoretical and Applied Genetics, 119(2), 353–369. 10.1007/s00122-009-1044-z

Peleg, Z., Saranga, Y., Yazici, A., Fahima, T., Ozturk, L., & Cakmak, I. (2008). Grain zinc, iron and protein concentrations and zinc-efficiency in wild emmer wheat under contrasting irrigation regimes. Plant and Soil, 306(1–2), 57–67. 10.1007/s11104-007-9417-z

Peterson, B. G., Carl, P., Boudt, K., Bennett, R., Ulrich, J., Zivot, E., Lestel, M., Balkissoon, K., & Wuertz, D. (2014). PerformanceAnalytics: Econometric tools for performance and risk analysis. R Package Version, 1(3).

Puranik, S., Sahu, P. P., Beynon, S., Srivastava, R. K., Sehgal, D., Ojulong, H., & Yadav, R. (2020). Genome-wide association mapping and comparative genomics identifies genomic regions governing grain nutritional traits in finger millet (*Eleusine coracana* L. Gaertn.). PLANTS, PEOPLE, PLANET, 2(6), 649–662. 10.1002/ppp3.10120

R Core Team, R. C. (2020). R: A Language and Environment for Statistical Computing.

Reda, A. (2014). Achieving Food Security in Ethiopia by Promoting Productivity of Future World Food Tef: A Review. Advances in Plants & Agriculture Research, 2(2). 10.15406/apar.2015.02.00045

Remington, D. L., Thornsberry, J. M., Matsuoka, Y., Wilson, L. M., Whitt, S. R., Doebley, J., Kresovich, S., Goodman, M. M., & Buckler, E. S. (2001). Structure of linkage disequilibrium and phenotypic associations in the maize genome. Proceedings of the National Academy of Sciences, 98(20), 11479–11484. 10.1073/pnas.201394398

Saturni, L., Ferretti, G., & Bacchetti, T. (2010). The Gluten-Free Diet: Safety and Nutritional Quality. Nutrients, 2(1), 16–34. 10.3390/nu2010016

Schmidt, S. B., Jensen, P. E., & Husted, S. (2016). Manganese Deficiency in Plants: The Impact on Photosystem II. Trends in Plant Science, 21(7), 622–632. 10.1016/j.tplants.2016.03.001

Sehgal, A., Sita, K., Siddique, K. H. M., Kumar, R., Bhogireddy, S., Varshney, R. K., HanumanthaRao, B., Nair, R. M., Prasad, P. V. V., & Nayyar, H. (2018). Drought or/and Heat-Stress Effects on Seed Filling in Food Crops: Impacts on Functional Biochemistry, Seed Yields, and Nutritional Quality. Frontiers in Plant Science, 9, 1705. 10.3389/fpls.2018.01705

Shumoy, H., Pattyn, S., & Raes, K. (2018). Tef protein: Solubility characterization, in-vitro digestibility and its suitability as a gluten free ingredient. LWT, 89, 697–703. 10.1016/j.lwt.2017.11.053

Siddique, K. H. M., Li, X., & Gruber, K. (2021). Rediscovering Asia’s forgotten crops to fight chronic and hidden hunger. Nature Plants, 7(2), 116–122. 10.1038/s41477-021-00850-z

Stagnari, F., Galieni, A., & Pisante, M. (2016). Drought stress effects on crop quality. Water Stress and Crop Plants: A Sustainable Approach, 2, 375–392.

Stangoulis, J., & Sison, C. (2008). Crop sampling protocols for micronutrient analysis. Harvest Plus Tech Monogr Ser, 7, 1–20.

Stevens, G. A., Beal, T., Mbuya, M. N. N., Luo, H., Neufeld, L. M., Addo, O. Y., Adu-Afarwuah, S., Alayón, S., Bhutta, Z., Brown, K. H., Jefferds, M. E., Engle-Stone, R., Fawzi, W., Hess, S. Y., Johnston, R., Katz, J., Krasevec, J., McDonald, C. M., Mei, Z., … Young, M. F. (2022). Micronutrient deficiencies among preschool-aged children and women of reproductive age worldwide: A pooled analysis of individual-level data from population-representative surveys. The Lancet Global Health, 10(11), e1590–e1599. 10.1016/S2214-109X(22)00367-9

Tadele, Z. (2019). Orphan crops: Their importance and the urgency of improvement. Planta, 250(3), 677–694. 10.1007/s00425-019-03210-6

Tao, H., Dittert, K., Zhang, L., Lin, S., Römheld, V., & Sattelmacher, B. (2007). Effects of soil water content on growth, tillering, and manganese uptake of lowland rice grown in the water-saving ground-cover rice-production system (GCRPS). Journal of Plant Nutrition and Soil Science, 170(1), 7–13. 10.1002/jpln.200625033

Tietel, Z., Simhon, E., Gashu, K., Ananth, D. A., Schwartz, B., Saranga, Y., & Yermiyahu, U. (2020). Nitrogen availability and genotype affect major nutritional quality parameters of tef grain grown under irrigation. Scientific Reports, 10(1), 14339. 10.1038/s41598-020-71299-x

Vadez, V., Grondin, A., Chenu, K., Henry, A., Laplaze, L., Millet, E. J., & Carminati, A. (2024). Crop traits and production under drought. Nature Reviews Earth & Environment, 5(3), 211–225. 10.1038/s43017-023-00514-w

VanBuren, R., Man Wai, C., Wang, X., Pardo, J., Yocca, A. E., Wang, H., Chaluvadi, S. R., Han, G., Bryant, D., Edger, P. P., Messing, J., Sorrells, M. E., Mockler, T. C., Bennetzen, J. L., & Michael, T. P. (2020). Exceptional subgenome stability and functional divergence in the allotetraploid Ethiopian cereal teff. Nature Communications, 11(1), 884. 10.1038/s41467-020-14724-z

VanRaden, P. M. (2008). Efficient Methods to Compute Genomic Predictions. Journal of Dairy Science, 91(11), 4414–4423. 10.3168/jds.2007-0980

Vavilov, N. I. Nikolaĭ I. (1951). The origin, variation, immunity and breeding of cultivated plants, Translated from the Russian by Chester KS, Ronald Press Co., New York. Ronald Press Co. https://cir.nii.ac.jp/crid/1130282269828906624

Villanueva, M., Abebe, W., Pérez-Quirce, S., & Ronda, F. (2022). Impact of the Variety of Tef [Eragrostis tef (Zucc.) Trotter] on Physical, Sensorial and Nutritional Properties of Gluten-Free Breads. Foods, 11(7), 1017. 10.3390/foods11071017

Von Berg, J., Ten Dam, M., Van Der Laan, S. W., & De Ridder, J. (2022). PolarMorphism enables discovery of shared genetic variants across multiple traits from GWAS summary statistics. Bioinformatics, 38(Supplement_1), i212–i219. 10.1093/bioinformatics/btac228

Wang, J., & Zhang, Z. (2021). GAPIT Version 3: Boosting Power and Accuracy for Genomic Association and Prediction. Genomics, Proteomics & Bioinformatics, 19(4), 629–640. 10.1016/j.gpb.2021.08.005

Wickham, H., Chang, W., & Wickham, M. H. (2016). Package ‘ggplot2.’ Create Elegant Data Visualisations Using the Grammar of Graphics. Version, 2(1), 1–189.

Woldeyohannes, A. B., Iohannes, S. D., Miculan, M., Caproni, L., Ahmed, J. S., De Sousa, K., Desta, E. A., Fadda, C., Pè, M. E., & Dell’Acqua, M. (2022). Data-driven, participatory characterization of farmer varieties discloses teff breeding potential under current and future climates. eLife, 11, e80009. 10.7554/eLife.80009

Xu, Z., Lai, T., Li, S., Si, D., Zhang, C., Cui, Z., & Chen, X. (2018). Promoting potassium allocation to stalk enhances stalk bending resistance of maize (Zea mays L.). Field Crops Research, 215, 200–206. 10.1016/j.fcr.2017.10.020

Yazici, M. A., Asif, M., Tutus, Y., Ortas, I., Ozturk, L., Lambers, H., & Cakmak, I. (2021). Reduced root mycorrhizal colonization as affected by phosphorus fertilization is responsible for high cadmium accumulation in wheat. Plant and Soil, 468(1–2), 19–35. 10.1007/s11104-021-05041-5

Zhu, F. (2018). Chemical composition and food uses of teff (Eragrostis tef). Food Chemistry, 239, 402–415. 10.1016/j.foodchem.2017.06.101

Zorb, C., Senbayram, M., & Peiter, E. (2014). Potassium in agriculture – Status and perspectives. Journal of Plant Physiology, 171(9), 656–669. 10.1016/j.jplph.2013.08.008

